# Context-dependent triggering of STING-interferon signaling by CD11b agonists supports anti-tumor immunity in mouse models and human cancer patients

**DOI:** 10.1101/2022.03.22.485233

**Authors:** Xiuting Liu, Graham D. Hogg, Chong Zuo, John M Baer, Varintra E. Lander, Liang Kang, Nicholas C. Borcherding, Brett L. Knolhoff, Robin E. Osterhout, Anna V. Galkin, Jean-Marie Bruey, Laura L. Carter, Cedric Mpoy, Julie K. Schwarz, Haeseong Park, Vineet Gupta, David G. DeNardo

**Author notes:** Corresponding author: David G. DeNardo, Department of Medicine, 425 South Euclid Ave, St. Louis, MO 63110.

## Abstract

Chronic activation of inflammatory pathways and suppressed interferon signaling are hallmarks of myeloid cells in immunosuppressive tumors that drive poor responsiveness to conventional and immune therapies. Previous studies have identified agonistic activation of the CD11b integrin as a potential strategy to enhance anti-tumor immunity. However, the mechanisms by which CD11b-agonism reprogram tumor immunity are poorly understood, and this may impair patient selection and identification of effective treatment combinations. Herein we used a combination of *in vitro* systems, animal models, and samples from first in human clinical trials of the CD11b-agonist GB1275 to identify the mechanism of action of this approach and identify combinations for further testing. We found that CD11b agonism altered tumor-associated macrophage (TAM) phenotypes by simultaneously repressing NFκB/IL1-signaling and activating interferon (IFN) gene expression. Repression of NFκB/IL-1 signaling was due to rapid degradation of p65 protein by the proteosome and was not context dependent. In contrast, CD11b agonism triggered mitochondrial dysfunction to stimulate STING-induced, STAT1-mediated interferon signaling. The magnitude of CD11b agonist induction of STING/IFN signaling was dependent on the tumor microenvironment and was significantly amplified by cytotoxic therapies. Using tissues from phase I clinical trials, we demonstrated that GB1275 treatment activated STING and STAT1 signaling in TAMs in human tumors. Together, these mechanisms allowed macrophages to augment anti-tumor T cell immunity. These studies identified potential mechanism-based therapeutic strategies for CD11b agonist use and identified potential patient populations more likely to benefit from them.

**Statement of significance:** CD11b agonists are a novel approach to reprogram myeloid cells in solid tumors. We show that GB1275, a CD11b-agonist, amplified STING/IFN signaling in TAMs to support anti-tumor immunity and this signaling is amplified further by cytotoxic therapy. These studies support new treatment strategies for advanced solid tumors with myeloid immunosuppression.

## Introduction

The presence of high numbers of tumor-associated macrophages (TAMs) has been associated with poor clinical outcomes in many cancer types (1). Even though multiple cancer models have demonstrated TAMs can foster tumor progression and immune suppression (2); TAMs also have considerable plasticity and can play both pro-tumoral and anti-tumoral roles (3). In addition, some solid tumors, such as pancreatic ductal adenocarcinoma (PDAC), are characterized by a dense desmoplastic stroma. This desmoplastic stroma includes elements of both fibrosis and abundant myeloid infiltration, which may synergize to drive immunosuppressive and wound healing programs (4–6). Thus, targeting the tumor microenvironment by re-shaping TAMs toward anti-tumor phenotypes is an attractive therapeutic strategy for advanced cancer patients (7).

TAMs are highly plastic and capable of regulating multiple aspects of tumor promotion and restraint (1,8,9). Anti-tumorigenic macrophages are characterized by high expressions of tumor necrosis factor (TNFα), IL12a, inducible nitric oxide synthase (iNOS) or MHC class II molecules, and T cell attracting chemokines, such as CXCL9 and CXCL10 (10). In contrast, pro-tumorigenic macrophages expression high levels of IL-10, the IL-1 decoy receptor, and IL-1Ra, arginase-1 and the scavenger receptors CD163, CD204, or CD206 (11, 12). In human PDAC, TAMs are abundant in the tumor tissues and high numbers corelate with poor clinical outcomes (13, 14). Considering the critical role of macrophages in the initiation and progression of many cancers, studies have examined the efficiency of reprogramming TAMs as an anti-cancer therapy. Some of the earliest approaches focused on TAM depletion. In preclinical models blocking key chemokine receptors (e.g., CCR2 or CXCR1/2) or impairing TAM survival by inhibiting colony stimulating factor-1 receptor (CSF1R) have shown efficacy in slowing progression and improving responses to a variety of agents (15–17). However, to date, these approaches, while tested, have yet to achieve clinically significant benefits in solid tumors (18). This is likely because these myeloid-ablative strategies are subject to significant compensatory actions by untargeted subsets of monocytes, granulocytes, and/or tissue resident macrophages that may ultimately limit their therapeutic efficacy in humans (19, 20). An alternative strategy is to reprogram TAMs to support tumor immunity.

Previous studies identified CD11b as a candidate for immunotherapy (21–23). CD11b is composed of the αM (ITGAM) and β2 (CD18) integrins and is widely expressed on multiple myeloid cell subsets. Agonism of CD11b by small molecules leads to a modestly reduced number of tumor-infiltrating immunosuppressive myeloid cells, such as TAMs, monocytes, and granulocytes, and a corresponding increase in tumor infiltrating T cells in PDACs and other mouse models (21–23). Additionally, agonism of CD11b has the potential to reprogram TAMs, including increased expression of IFN-related genes (21). These data also led to early-phase clinical testing in solid tumors (NTC04060342). In this paper, we investigated the cellular and molecular mechanism by which CD11b agonists induce anti-tumor immunity and identify combination therapies to optimize this.

## Results

### Cancer-specific differences in tumor immunity pathways in TAMs

The TME of PDACs has been proposed to be immunosuppressive and shown to be heavily infiltrated by myeloid cells (24–26). As such, it has been proposed that TAMs and other myeloid cells are key drivers of immune suppression in PDACs (27–29). However, if PDAC TAMs have unique immunosuppressive and tissue remodeling phenotypes, compared to TAMs in other major cancers, is not known. To begin to understand the PDAC unique pathways that may impair tumor immunity, we combined publicly available single-cell RNA-seq (scRNAseq) data sets across ten cancer types to identify greater than 54,000 TAMs (Fig. 1A). As expected, even after significant integration, Uniform Manifold Approximation and Projection (UMAP) analysis still shows some segregation of TAMs by tumor types, suggesting that the tumor-specific phenotypes and transcriptional features (Fig. 1A, Sup. Fig. 1A). To compare TAMs across cancers, we used gene set enrichment analysis (GSEA). We found that pathway specific gene sets formed unique patterns for each cancer type, again suggesting cancer-specific phenotype bias in TAMs. Comparing PDAC TAMs to TAMs in other major cancers, we observed some of the highest levels of the TGFβ, WNT, NFκB and IL4/IL13 pathways and high expression of hypoxia and glycolysis gene sets (Fig. 1B, Sup. Fig. 1B-C). These results may be indicative of the highly fibroinflammatory TME that is a hallmark of PDAC. We also observed strong congruence for these pathways seen across different GSEA databases, including Hallmark, PID or Reactome (Sup. Fig. 1B-C). By contrast, we observed that PDAC TAMs had some of the lowest levels of type-1 interferon-signaling, antigen presentation and reactive oxygen species, oxidative phosphorylation, and peroxisome gene sets (Fig. 1B, Sup. Fig. 1B-C). Taken together, these data were consistent with what would be predicted for a highly desmoplastic cancer, like PDAC, and suggested that in such cancers, augmenting interferon signaling may be key to tumor immunity in this disease.

**Figure 1.**
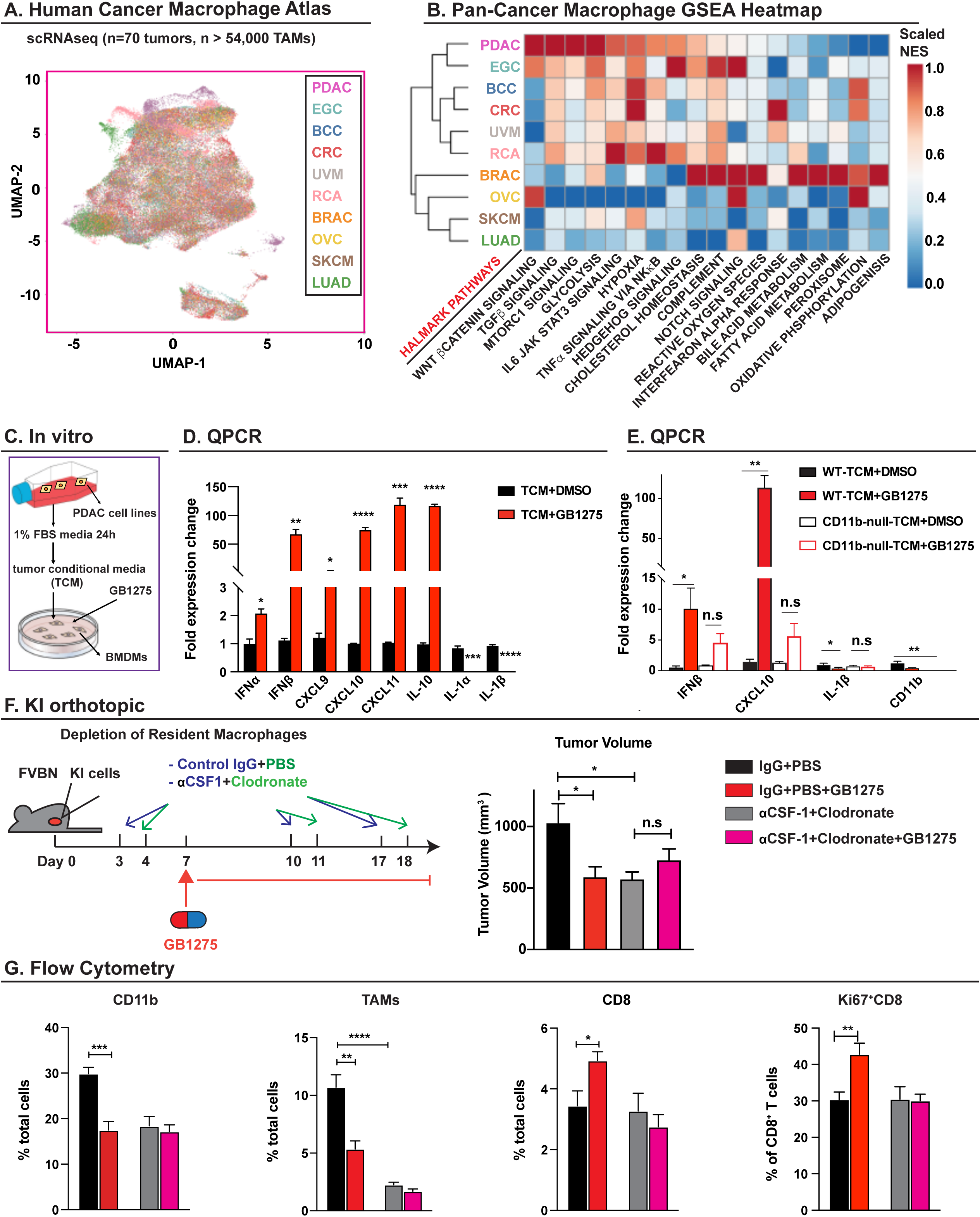
CD11b agonist anti-tumor activity is dependent on tumor-associated macrophage (TAM)s reprograming. **A**. UMAP visualization of over 54000 tumor-associated macrophages (TAMs) from different cancer types, including pancreatic cancer (PDAC), early gastric cancer (EGC), Basal cell carcinoma (BCC), colorectal cancer (CRC), uveal melanoma (UVM), Renal cancer (RCA), breast cancer (BRAC), ovarian serous cystadenocarcinoma (OVC), skin cutaneous melanoma (SKCM) and lung adenocarcinoma (LUAD) analyzed by scRNA-seq and integrated across 70 tumors. Clusters were annotated for cancer types. **B**. Gene set enrichment analysis (GSEA) identified pathway enrichment in TAMs from above cancer types. **C**. The in vitro model used to analyze the regulation by GB1275 on macrophages. Bone marrow-derived macrophages (BMDMs) were treated with vehicle or GB1275 in the presence or absence of pancreatic ductal adenocarcinoma (PDAC)-conditioned media for 7 h. **D**. Quantitative-PCR mRNA expression analysis of BMDMs treated with PDAC-conditioned media ± vehicle or GB1275 for 7 h. Changes in gene expression are depicted as the fold change from the vehicle baseline. **E**. BMDMs isolated from wild-type and CD11b-null mice were treated by the above stimulus. Changes in gene expression are depicted as the fold change from the vehicle baseline. **F.** Mice bearing syngeneic orthotopic KI tumors with three doses of IgG+PBS/ aCSF-1+Clodronate. On day 7, mice were treated with vehicle or GB1275 (120mg/kg) for 12 days. Tumor volume at day 12 of treatment is depicted (n = 7–9/group). **G**. Frequencies of tumor infiltrating CD11b^+^ cells, tumor-associated macrophages (TAMs), CD8^+^T cells, and Ki67^+^ CD8^+^T cells in the above orthotopic KI PDAC tissues (n=6/group). All graphs depict the mean ± SEM and *denotes a value of p < 0.05 using the two-sided *t*-test or analysis of variance. *In vitro* data are representative of three independent and consistent experiments.

### CD11b agonist anti-tumor activity is dependent on TAM reprograming

CD11b integrins can be critical for myeloid recruitment into the TME, including regulation of adhesion, migration, and chemotaxis of myeloid cells (21). But CD11b integrins can also regulate myeloid cell phenotypes. In previous studies, we used CD11b agonists to target myeloid cells, and found it led to reduced tumor infiltrating myeloid cell numbers, restraint of tumor progression through enhancement of tumor immunity (22). These small molecule agonists were formally named either LA1 or ADH503, but will be referred to as GB1275 herein, given their current clinical format (Sup Fig. 2A and NTC04060342). As in prior studies, we found macrophages polarized with PDAC tumor cell conditioned media (TCM) and treated *in vitro* with GB1275 directly upregulated multiple interferon (IFN)-regulated genes, but downregulated IL-1 related genes (Fig. 1C-D). These effects were not seen in macrophages derived from CD11b-null mice (Fig. 1E). While these data were consistent with prior studies, it remained unclear whether reduced myeloid cell number or TAM reprograming are the key drivers of the efficacy observed with CD11b-agonists.

To begin to study this, we first confirmed that the GB1275 formulation exhibited anti-tumor activity *in vivo* through CD11b activation. We used syngeneic cells derived from the genetic KPC (p48-Cre;LSL-Kras^G12D^;Trp53^flox/+^) mouse PDAC model, named KP2 (30). KP2 PDAC cells were orthotopically implanted in both wild-type (WT) and CD11b-null mice. As expected, GB1275 repressed tumor progression (Sup. Fig. 2B), decreased CD11b^+^ myeloid cell infiltration (Sup. Fig. 2C, 3A), and increased CD8^+^ T cell recruitment and proliferation in PDAC tissues (Sup. Fig. 2D, 3B). However, these effects of GB1275 were lost in CD11b-null mice (Sup. Fig. 2B-D). As CD11b agonists could directly alter macrophage phenotype, we next sought to determine if TAMs were necessary for efficacy and induction of anti-tumor immunity. We depleted tissue macrophages using a combination of anti-CSF-1 IgG and liposomal clodronate (αCSF1/CLD), as previously published (31). Mice bearing established syngeneic orthotopic tumors derived from KrasG12D-Ink4a^f/f^ (KI cells) were treated with αCSF1/CLD and analyzed by flow cytometry. As expected αCSF1/CLD depleted >80% of TAMs, which led to reduced tumor burden on its own but did not increase T cell infiltration (Fig. 1F, G). As previously, GB1275 treatment reduced KI tumor progression and increased CD8^+^ CTL infiltration and proliferation; however, these changes were dependent on the presence of TAMs (Fig. 1F, G). These data suggested that GB1275 activated CD11b integrins on TAMs to drive the induction of anti-tumor immunity and tumor restraint.

### CD11b activation on TAMs results in proliferation of effector T cells

Our prior studies have shown that the anti-tumor activity of CD11b-agonists was dependent on T cell immunity (22). Notably, T cells did not express high levels of CD11b, so we next aimed to understand how CD11b agonist myeloid reprograming changed T cell phenotypes. To accomplish this objective, mass cytometry (CyTOF) was employed on orthotopic KI tumors treated with vehicle or GB1275. After 12 days of treatment, GB1275 decreased the tumor burden (Fig. 2A). As expected, we observed decreased infiltration of TAMs, and increased CD8^+^ T cell numbers and proliferation by flow cytometry (Fig. 2B). UMAP analysis of CyTOF data on these same tissues identified five subsets from CD8^+^ T cells, five subsets of CD4^+^ affected T cells, and two subsets of Tregs (Fig. 2C and Sup. Fig. 4A). Across all CD8^+^ T cell subsets, we observed increased expressions of granzyme B (GZB) and Ki67 (Fig. 2D). Analyzing changes in CD8^+^ T cell subsets, we found a significant increase of proliferative effector CD8^+^ T cells (cluster 5), which expressed high levels of CD44, GZB, Tbet, and Ki67 (Fig. 2E). In contrast, GB1275 treatment led to decreases in CD103^+^ resident memory (cluster 2) and activated non-proliferative non-effector CD8^+^ T cells (cluster 3), which expressed CD39, CD44 and Tbet, but no GZB or Ki67 (Fig. 2E). Notably, across all subsets, we found a trend towards increased PD-1, CTLA-4, and TIM3 expressions with treatment in this model (Sup. Fig. 4B). These data demonstrated that CD11b agonism-induced changes in myeloid cells drove CTLs toward proliferative effector phenotypes.

**Figure 2.**
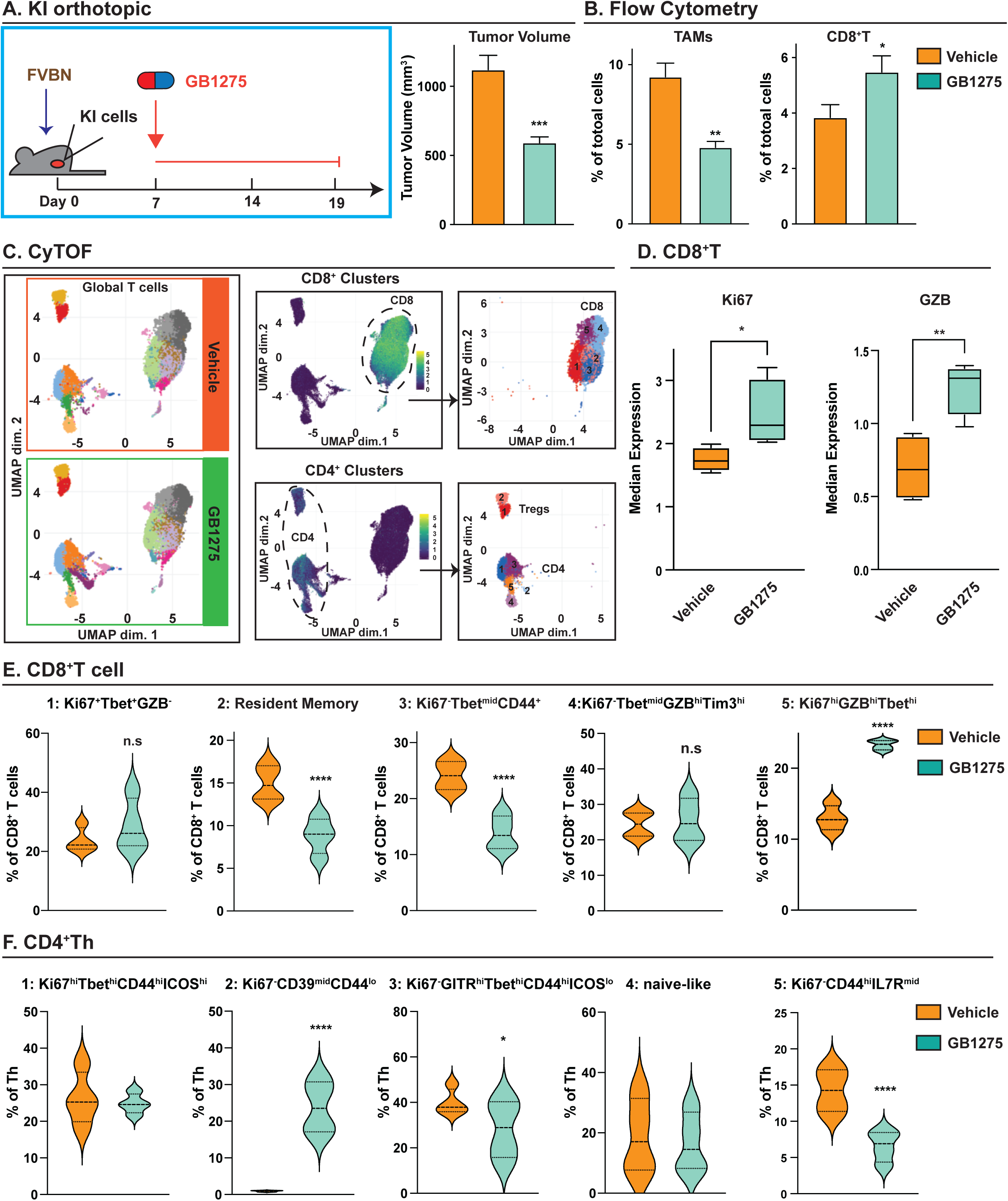
CD11b activation on tumor-associated macrophages results in more proliferative effector T cells. **A**. Syngeneic orthotopic model of KI cells treated with vehicle or GB1275 (60mg/kg) for 12 days (left). (Right) Tumor burden at 12-day treatment from the above two groups shown by tumor volume (n = 8/group). **B**. Relative frequencies of tumor-infiltrating macrophages and CD8^+^ T cells from the above mice (n = 7/group). **C**. CyTOF UMAP plot of tumor infiltrating T cells (left). Subpopulations of CD8^+^ T cell (upper) and CD4^+^ T cells, including T^Regs^ (bottom). **D**. Median expressions of Ki67 (left) and Granzyme B (right) in CD8^+^ T cells. **E**. Percentage of individual subclusters in CD8^+^T cells. **F**. Percentage of individual subclusters in CD4^+^T helper cells. Graphs depict the mean ± SEM and *denotes a value of p < 0.05 using a two-sided *t*-test.

When analyzing CD4^+^ T cells, GB1275 exhibited more impact on T helper cell (Th) phenotypes compared to FOXP3^+^ regulatory T cell (T_Regs_) subsets. When analyzing Th subclusters, we found that GB1275 decreased memory CD4^+^ T cells (cluster 5), characterized by high expressions of CD44 and IL-7R, and cluster 3, expressing high GITR, Tbet, CD44, and low ICOS and Ki67, indicative of non-proliferative activated CD4^+^ T cells. The proliferative activated Th (cluster 1) was not changed in the GB1275-treated group, when compared with the vehicle. In contrast, GB1275 induced the accumulation of cluster 2, with low expression of CD39 and CD44 and possibly early effector CD4^+^ T cells (Fig. 2F). In addition, the naïve-CD4^+^ T cells (cluster 4) and two subpopulations of Tregs were not changed in frequency by GB1275 (Sup Fig. 4C). From abundance expression analyses, neither exhaustion markers/checkpoints nor proliferation markers, including PD-1, CTLA-4, TIM3, and Ki67, was affected by GB1275 on CD4^+^ T cells (Sup Fig. 4D, E). These data suggest that GB1275 anti-tumor activity in the PDAC models is predominantly driven by modulation of TAM mediated recruitment of the proliferative effector CD8^+^ T cell subset.

### NFκB/IL-1 signaling in TAMs is downregulated by agonism of CD11b

To understand how CD11b-agonism might induce changes in TAMs to support T cell effector function, we next studied how CD11b agonists changed TAM phenotype in-vivo. To accomplish this, we performed scRNAseq on CD45^+^ cells isolated from KP2 PDAC tissue from mice treated with vehicle or GB1275. UMAP analysis distinguished 21 clusters in CD45^+^ cells (Sup. Fig. 5A). The TAM/monocyte populations were isolated and re-clustered (Fig. 3A) to identify three subpopulations of TAMs and monocytes cluster, denoted as TAM1, TAM2, TAM3, and Mo (Fig. 3A, Sup. Fig. 5B). Notably, we observed only a modest shift in TAM subclusters in GB1275-treated mice (Fig. 3A), suggesting that TAM subsets or origins were not as markedly changed (1, 31). Interestingly, as above, the number of TAMs in PDAC tissues were uniformly decreased in this experiment by GB1275, suggesting that decreases occur across all subsets. Notably, we did observe significant changes in TAM phenotypes across all these subsets. To better understand this, we identified differentially-expressed genes (DEGs) and conducted gene set enrichment analysis (GSEA) on total TAMs. We observed enrichment in genes in several integrin pathways in GB1275-treated samples, consistent with activation of the CD11b integrins, αM/β2 (Fig. 3B). Additionally, we observed that oxidative phosphorylation and reactive oxygen species (ROS) signatures were increased after GB1275 treatment, indicating possible oxidative stress in TAMs. Notably, inflammatory signaling, like TNFα and NFκB signaling, was downregulated after CD11b agonism, and the majority of NFκB/IL1 signaling pathway-associated genes were significantly decreased in TAMs from GB1275-treated mice (Fig. 3B-D). These included the IL1- related genes, IL-1β, IL1R1, and CCL2 (Fig. 3D). Together these data suggest that CD11b-agonists altered TAM phenotypes *in vivo*, but the mechanisms at play are not understood.

**Figure 3.**
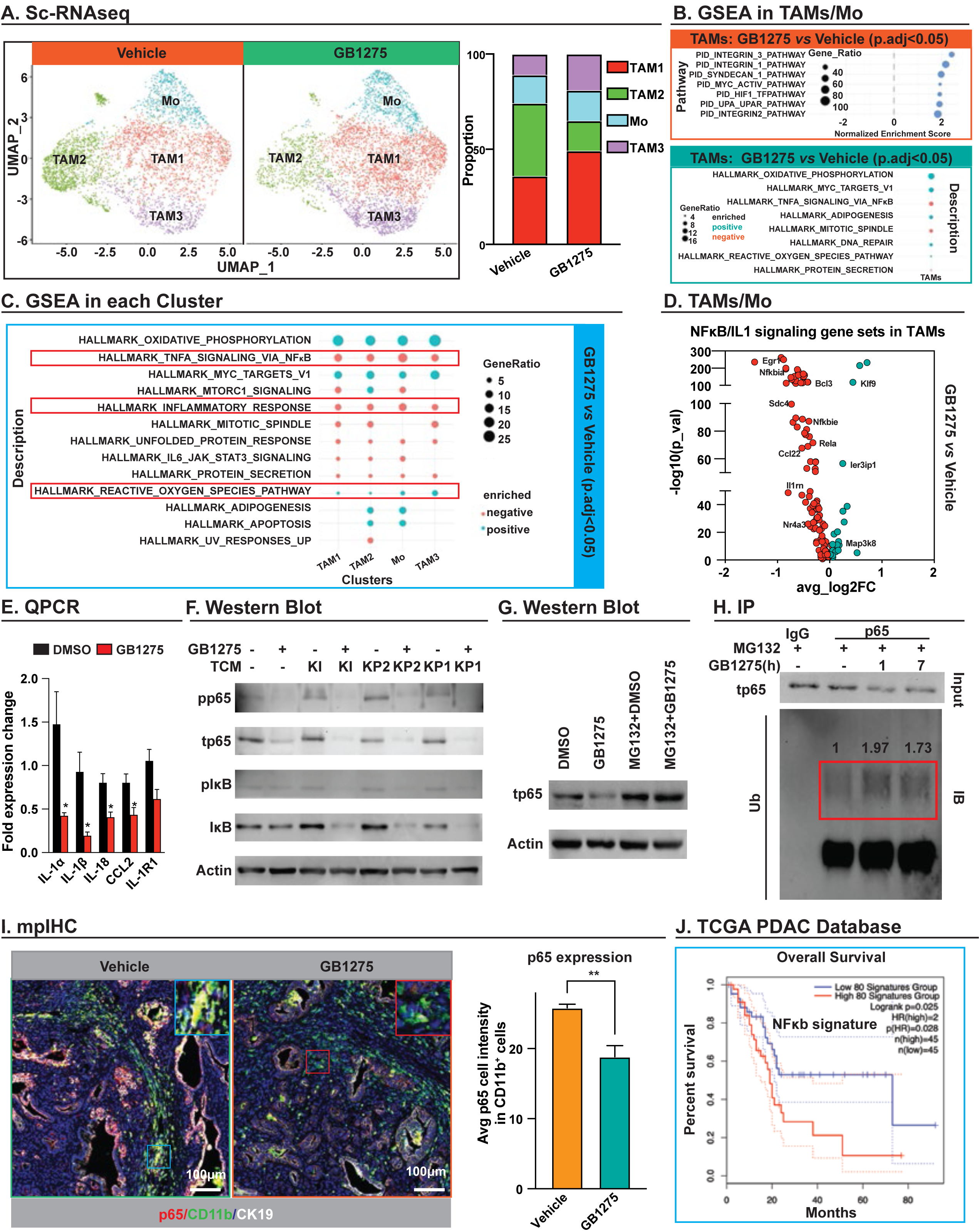
NFκB/IL-1 signaling in tumor-associated macrophages (TAMs) is downregulated by CD11b activation. **A**. UMAP scRNAseq plots show the subclusters of TAM/monocyte population in vehicle and 14-day GB1275 (120mg/kg)-treated KP2 syngeneic model (left). Percentages of cells in individual clusters in the TAM/monocytes population by their conditions (right). **B**. Gene set enrichment analysis (GSEA) identified pathway enrichment in whole TAM/monocyte populations by their conditions. **C**. GSEA identified pathway enrichment in four subclusters of a TAM/monocyte population by their conditions. **D**. Volcano plot depicting differentially-expressed genes within the NFκB/IL-1 signaling pathway from the whole TAM/monocyte population in the GB1275-treated group, when compared with the vehicle with significant difference. **E**. Quantitative PCR mRNA expression analysis of bone marrow-derived macrophages (BMDMs) treated by GB1275 for 7 h. Changes in gene expression are depicted as the fold change from the vehicle baseline. **F**. Representative immunoblot for pp65, total p65, pIκB, total IκB, and β-actin in BMDMs treated with GB1275 ± TCM for 7 h. **G**. BMDMs were pretreated with MG132 (10uM) 1 hour earlier, and then treated with GB1275 for seven hours. Representative immunoblot of total p65 and β-actin (loading control). **H**. BMDMs were pretreated with MG132 (10 µM) 1 hour earlier, and then treated with GB1275 for one or seven hours. The p65 purified protein was isolated by p65 antibody. Immunoblot for p65 from total lysates and polymer-ubiquitin from purified protein showed the combination of ubiquitin with p65. **I**. Representative mpIHC staining for p65, CD11b and tumor marker CK19 in tumors from 14-day vehicle and GB1275 (120mg/kg)- treated KPPC mice. Scale bar, 100 μm. Right, average p65 cell intensity in CD11b^+^ cells from each treatment group (n = 6−7 mice per group). **J**. Kaplan-Meier survival curves for top 80 downregulated NFκB/ IL-1 signaling-related genes from scRNAseq (D) in TCGA patient dataset for pancreatic adenocarcinoma (PAAD). Graphs show the mean ± SEM; *denotes p < 0.05 using the two-sided *t*-test or log-rank test. All, *in vitro* data are representative of three independent repeat experiments.

Consistent with our observations *in vivo*, *in vitro* CD11b agonism decreased expression of IL-1 signaling-associated genes in macrophages. Macrophages treated for 7 h with GB1275 had decreased IL-1α, IL-1β, IL-18, and CCL2 gene expression (Fig. 3E). Decreases in IL-1 genes by GB1275 were dependent on CD11b, but independent of the presence or sources of tumor-conditioned media (TCM; Sup. Fig. 5C-F). We also observed downregulation of IL-1β in human monocyte-derived macrophage cultures, suggesting species conservation (Sup. Fig. 5E). Because scRNAseq data suggested downregulation of NFκB and inflammatory signaling by GB1275, we assessed these pathways in bone marrow-derived macrophages (BMDMs) under standard CSF1 culture conditions or under “tumor-polarizing” conditions by adding TCM. We found that under both conditions, GB1275 decreased both total and phosphorylated NFκB p65 and IκB protein levels (Fig. 3F). Notably, GB1275’s ability to decrease NFκB p65 expression was independent of the source or presence of TCM and was a dose-dependent effect (Fig. 3F; Sup. Fig. 6A). We analyzed several other inflammatory-related proteins and found that GB1275 also decreased phosphorylation of p38 MAPK and MK2 (Sup. Fig. 6A). Due to its known effect on IL-1 transcription (32, 33), we focused on CD11b-agonist regulation of NFκB. First, we found that GB1275 lost its ability to decrease IL-1 expression further when NFκB p65 was silenced by siRNA (Sup Fig. 6B, C). Next, we observed no decrease in NFκB p65 mRNA following GB1275 treatment (Sup Fig. 6D), suggesting that the regulation was likely centered on protein stability. Finally, we pretreated macrophages with MG132, a proteosome inhibitor, and found that GB1275 no longer decreased p65 protein expression (Fig. 3G), and we also detected that GB1275 increased poly-ubiquitination of p65 (Fig. 3H), which contributes to protostome reorganization and degradation (34, 35). Together these data suggest that CD11b agonism alters p65 degradation to regulate IL-1 expression.

We next sought to verify that CD11b-agonism decreased NFκB p65 in TAMs *in vivo*. To accomplish this, we treated p48-Cre;LSL-Kras^G12D^;Trp53^flox/flox^ (KPPC) mice with vehicle or GB1275 for 14 days and harvested pancreatic tumor tissues for multiplex immunohistochemistry (mpIHC) analysis. We serially stained for NFκB p65, CD11b, and CK19 (markers of PDAC cells) and found that GB1275-treated mice had reduced absolute numbers of p65^+^ CD11b^+^ myeloid cells (Sup Fig. 6E) and reduced p65 protein expression levels in the remaining CD11b^+^ myeloid cells (Fig. 3I). Based on downregulated NFκB/IL-1 signaling by GB1275 in our study, we next analyzed whether this signaling downregulation was related to favorable clinical outcomes in PDAC patients. To accomplish this aim, we identified genes from the Hallmark NFκB signaling pathway that were downregulated in TAMs following GB1275 treatment. We then used this CD11b-agonist specific gene signature to segregate PDAC patients in TCGA dataset and found that genes changed by CD11b-agonism were indicative of better overall survival in PDAC patients (Fig. 3J). These data suggested that CD11b agonist suppressed IL-1 signaling via NFκB degradation in TAMs, and that CD11b-agonist regulation of NFκB/IL-1 signaling might lead to favorable clinical outcomes in PDAC patients.

### CD11b activation of TAMs increases IFN/CXCL transcription via STING

As shown above, CD11b-agonists also upregulated IFN-related genes in TAMs, including CXCL9, 10, 11 and IFNα, and IFNβ (Fig. 1D). However, unlike IL-1-related genes, we found that GB1275’s ability to regulate IFN genes was not dependent on p65 expression (Sup. Fig. 6F). These data suggested an independent pathway of regulation. Notably, the induction of IFN genes by GB1275 was dependent on CD11b expression, and we only observed large changes of IFN genes in TAMs under TCM polarizing conditions or in co-cultured tumor cells (Fig. 4A; Sup. Fig. 7A-B). These data suggested that transcriptional changes in IFN genes might require tumor-specific cues in addition to activation of signaling using CD11b integrins. To investigate the potential signaling pathways involved, we performed reversed-phase protein arrays (RPPA) on BMDMs treated with GB1275 in the presence and absence of TCM. Compared with the vehicle, greater than 40 proteins were changed in BMDMs after either 4 or 7 h of GB1275 exposure in the context of TCM (Sup Fig. 7C). Among all the proteins, STING expression was increased by GB1275 alone, and further increased when GB1275 was added to BMDMs under TCM polarizing conditions (Fig. 4B). While activation of STING could explain both the induction of IFN genes *in vitro* and tumor immunity *in vivo*, the mechanisms by which CD11b would regulate STING were unknown.

**Figure 4.**
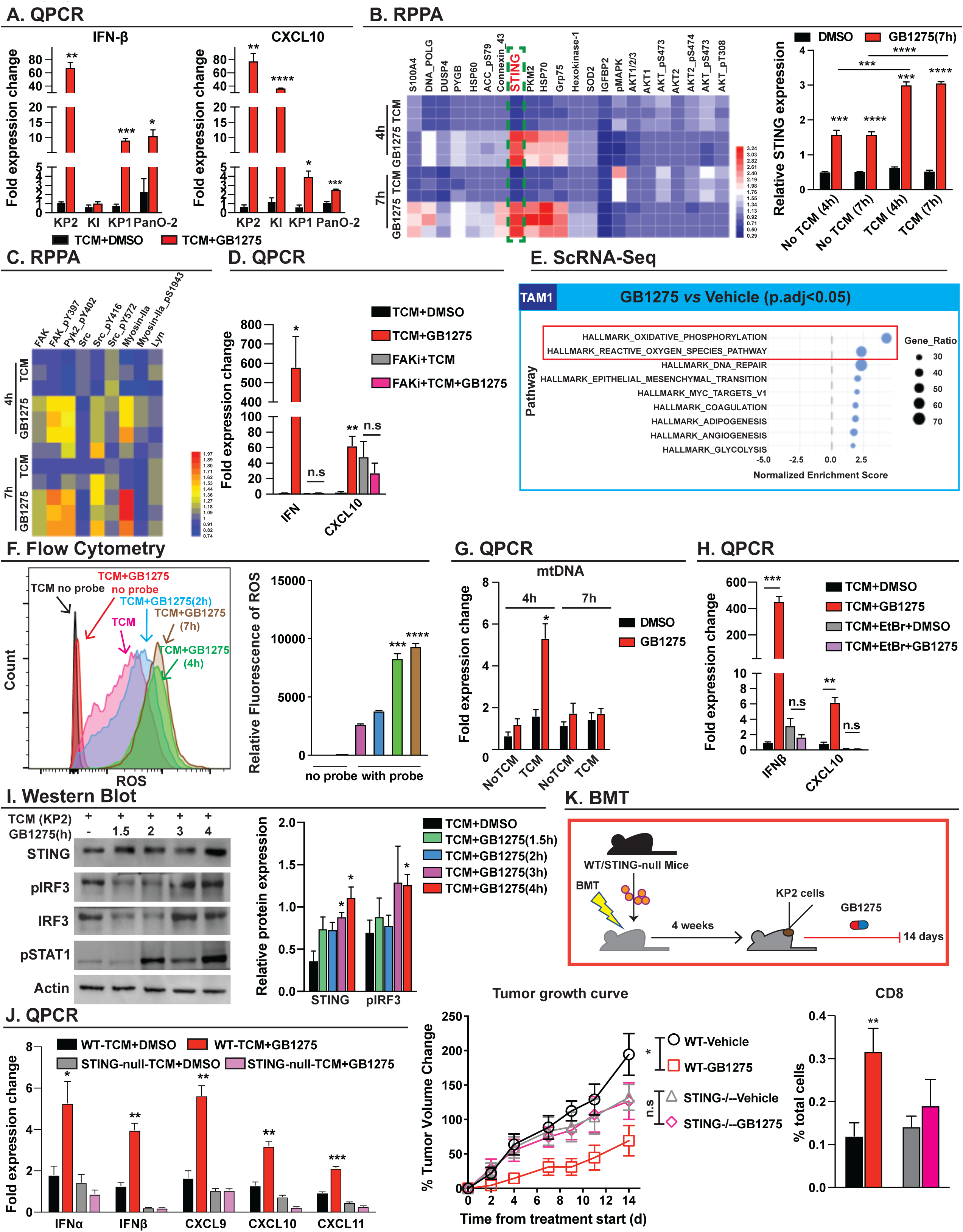
CD11b activation on TAMs increases IFN/CXCL transcription via STING. **A**. Quantitative PCR mRNA expression analysis of bone marrow-derived macrophages (BMDMs) treated with different pancreatic ductal adenocarcinoma (PDAC)-conditioned media ± vehicle or GB1275 for 7 h. Changes in gene expressions are depicted as the fold change from the vehicle baseline. **B**. Heat map of multiple protein relative expressions in BMDMs exposed to PDAC-conditioned media ± vehicle or GB1275 for 4 and 7 h (left). Relative STING expression after the above treatment (right). **C**. Heat map of relative expressions of FAK, pFAK, Src, and so on in BMDMs exposed to PDAC-conditioned media ± vehicle or GB1275 for 4 and 7 h. **D**. BMDMs were pretreated with FAK inhibitor (0.5 μM) 1 h ahead of TCM ± vehicle or GB1275 for 7 h. Quantitative PCR mRNA expression analysis of the above groups. Changes in gene expressions are depicted as the fold change from the vehicle baseline. **E**. Gene set enrichment analysis identified pathway enrichment in cluster TAM1 using scRNAseq in Fig. 3. **F**. BMDMs were stimulated by PDAC-conditioned media+GB1275 for different time points, and the intracellular levels of total reactive oxygen species (ROS) was measured by flow cytometry. Quantification of ROS production at indicated time points (right). **G**. Cytochrome C released in cytoplasm was detected by quantitative PCR analysis from BMDMs treated with tumor-conditioned medium (TCM) ± vehicle or GB1275 for both 4 and 7 h. Changes in gene expressions are depicted as the fold change from the vehicle baseline. **H**. BMDMs were pretreated with ethidium bromide (1.5 mg/mL) 1 h ahead of TCM ± vehicle or GB1275 for 7 h. Quantitative PCR mRNA expression analysis of the above cells with different stimulations. Changes in gene expressions are depicted as the fold change from the vehicle baseline. **I**. Representative immunoblot for STING, total IRF3, pIRF3, pSTAT1, and β-actin (loading control) in BMDMs treated with PDAC-conditioned media ± GB1275 for the indicated time points (left). Quantification of STING and pIRF3 relative expressions (right). **J**. Quantitative PCR mRNA expression analysis of BMDMs isolated from both wild-type and STING-null mice treated with PDAC-conditioned media ± vehicle or GB1275 for 7 h. Changes in gene expressions are depicted as the fold change from the vehicle baseline. **K**. C57/BL-6 mice were lethally irradiated and adoptively transferred with bone marrow cells from either wild-type mice or STING-null mice. After 4 weeks, these mice were adopted to establish a KP2-syngeneic model and after 10 days mice were treated with vehicle or GB1275 (120mg/kg) for 14 days (upper). Tumor growth curve from the above four groups expressed by percentages of tumor volume changes (n = 7−8/group) (left). Relative frequencies of tumor-infiltration CD8^+^ T cells from the above mice (n = 5−6/group). Graphs show the mean ± SEM; *denotes p < 0.05 using the two-sided *t*-test or analysis of variance. *In vitro* data are representative of three independent experiments.

As expected for an integrin agonist, proteomics analyses showed that FAK, PyK2, and SRC became phosphorylated, and expression of Myosin-IIa was increased in BMMACs after 4 hours of GB1275 treatment (Fig. 4C; Sup. Fig. 7D). GB1275-induced increases in IFN-β and CXCL10 mRNAs were reversed by FAK inhibition, suggesting that FAK activation was required for CD11b agonists to increase IFN transcription (Fig. 4D). ScRNAseq analysis of *in vivo* TAMs suggested that CD11b agonism activated oxidative phosphorylation and ROS pathways (Fig. 3B, C; Fig. 4E), which agreed with the observed increases in HSP70 and SOD proteins in *in vitro* macrophages by RPPA (Fig. 4B)(36, 37). These data suggested that CD11b agonists might induce oxidative stress in TAMs. Evaluating this hypothesis *in vitro*, we found GB1275 cooperated with TCM to enhance ROS production in a time-dependent manner and in a FAK activation-dependent manner (Fig. 4F; Sup. Fig. 7E). Next, we found that pretreatment of ROS scavengers ablated the induction IFN-β and CXCL10 transcriptions by GB1275 (Sup. Fig. 7F). Previous studies have shown that high ROS could be indicative of mitochondrial dysfunction and release of mtDNA, which can act as a primer for the cGAS-STING pathway (38). Indeed, we observed more mtDNA (cytochrome C) released into the cytosol following GB1275 combined with TCM treatment (Fig. 4G). Next, we depleted mtDNA from BMDMs by ethidium bromide (EtBr) treatment (39), and found the GB1275 induction of IFNβ and CXCL10 genes were markedly attenuated (Fig. 4H). Finally, immunoblot analysis of proteins from macrophages treated with GB1275 in TCM conditions validated the observed increased expressions of STING, and also showed that CD11b agonists induced increased phosphorylation of IRF3 and STAT1 (Fig. 4I). To determine if STING was responsible for CD11b agonist induction of IFN/CXCL genes, we silenced STING in BMDMs by siRNA using BMDMs from STING-null mice (Fig. 4J; Sup. Fig. 7G). In both conditions, GB1275 was unable to elevate IFN/CXCL mRNA levels (Fig. 4J; Sup. Fig. 7H). Notably, we also observed that GB1275 downregulation of IL1 mRNAs was independent of FAK activation, ROS, or STING expression (Sup. Fig. 8A-C), suggesting that NFkB p65 /IL-1 and STING/IFN were two separate signaling pathways under the regulation of CD11b activation. Together these data indicated that CD11b agonism led to ROS production and mitochondrial dysfunction, which facilitated STING/IFN signaling in “tumor-primed” macrophages *in vitro*.

We next sought to determine the importance of STING in CD11b agonist anti-tumor effects *in vivo*. To accomplish this, we transplanted mice with bone marrow (BM) from control or STING-null mice. After reconstitution we established syngeneic KP2 PDAC tumors and found that GB1275 treatment still restrained tumor progression and increased CD8^+^ T cell infiltration and proliferation in mice with WT BM, but these effects were lost in mice bearing STING-null BM (Fig. 4K; Sup. Fig. 7I). Taken together, these data suggested that CD11b agonists promoted tumor immunity through STING activation in myeloid cells.

We previously showed that CD11b agonist-treated tumors had fewer TAMs, and that CD11b agonists partially blocked monocyte trafficking (22). However, several groups have suggested that pancreatic tissue-resident macrophages may also expand by proliferation to sustain TAM numbers, with only modest contributions from trafficking monocytes (31,40,41). Interestingly, in these studies, we observed that the number of TAM in tumor tissues was not decreased by GB1275 in mice receiving STING-null BM (Sup Fig. 7I). These data suggest that STING activation may play a role in TAM turnover. Previous studies have established that induction of STING and IFN signaling increased macrophage turnover (42, 43). Thus, these data suggested that CD11b agonist activation of STING may contribute to reducing TAM number. Consistent with this possibility, *in vitro*, we observed that BMDMs had reduced cell numbers following prolonged high dose of 10 µM GB1275 exposure, and that this effect was partly dependent on STING expression (Sup. Fig. 8D). We also observed that prolonged GB1275 treatment led to reduced phosphorylated AKT (Sup. Fig. 8F) and higher levels of cleaved caspase 3 (Sup. Fig. 8E). These effects appeared to be specific to TAMs, as decreased granulocyte infiltration in GB1275-treated tumors did not require STING expression (Sup Fig. 7I). Taken together, these data suggested that CD11b agonists induced STING-mediated IFN signaling, which led to both tumor immunity and eventually TAM turnover.

### STING levels in TAMs in human PDAC correlates with tumor immunity and patient survival

We next sought to understand if we could use STING/IFN activation in TAMs as a prognostic biomarker for human cancers. To accomplish this, we first determined if we could observe STING activation *in vivo* in PDAC GEMMs following GB1275 exposure. In agreement with the *in vitro* data, KPC GEMMs treated with GB1275 had increased numbers of STING^+^ CD11b^+^ myeloid cells and higher STING protein expressions in CD11b^+^ myeloid cells compared to vehicle-treated mice (Fig. 5A, Sup. Fig. 9A). We next sought to understand the importance STING expression in TAMs in human PDAC cancer. To accomplish this aim, we used tissue microarrays (TMAs) consisting of PDAC tissues from 173 surgical patients. TMAs were stained by mpIHC for CD8α, STING, CD163, CD11b, and CK19 (Sup. Fig. 9C). When defining macrophages as CD11b^+^, CD163^+^, and CK19^-^, we found that STING^+^ TAMs ranged from 1−60% of total TAMs, with an average of 16% of total TAMs (Sup. Fig. 9B). In PDAC tissues higher percentages of STING^+^ TAMs (Fig. 5C) or STING^+^CD11b^+^ cells (Sup. Fig. 9B) correlated with increased CD8^+^ T cell infiltration. Furthermore, while previous studies have shown that high numbers of total CD163^+^ TAMs were correlated with poor clinical outcomes (44), we found patients whose PDAC had higher STING^+^ TAM infiltration exhibited longer overall survival (Fig. 5C). Together, these data suggested that activation of STING expression in TAMs might drive better T cell responses and improve clinical outcomes in human PDAC patients.

**Figure 5.**
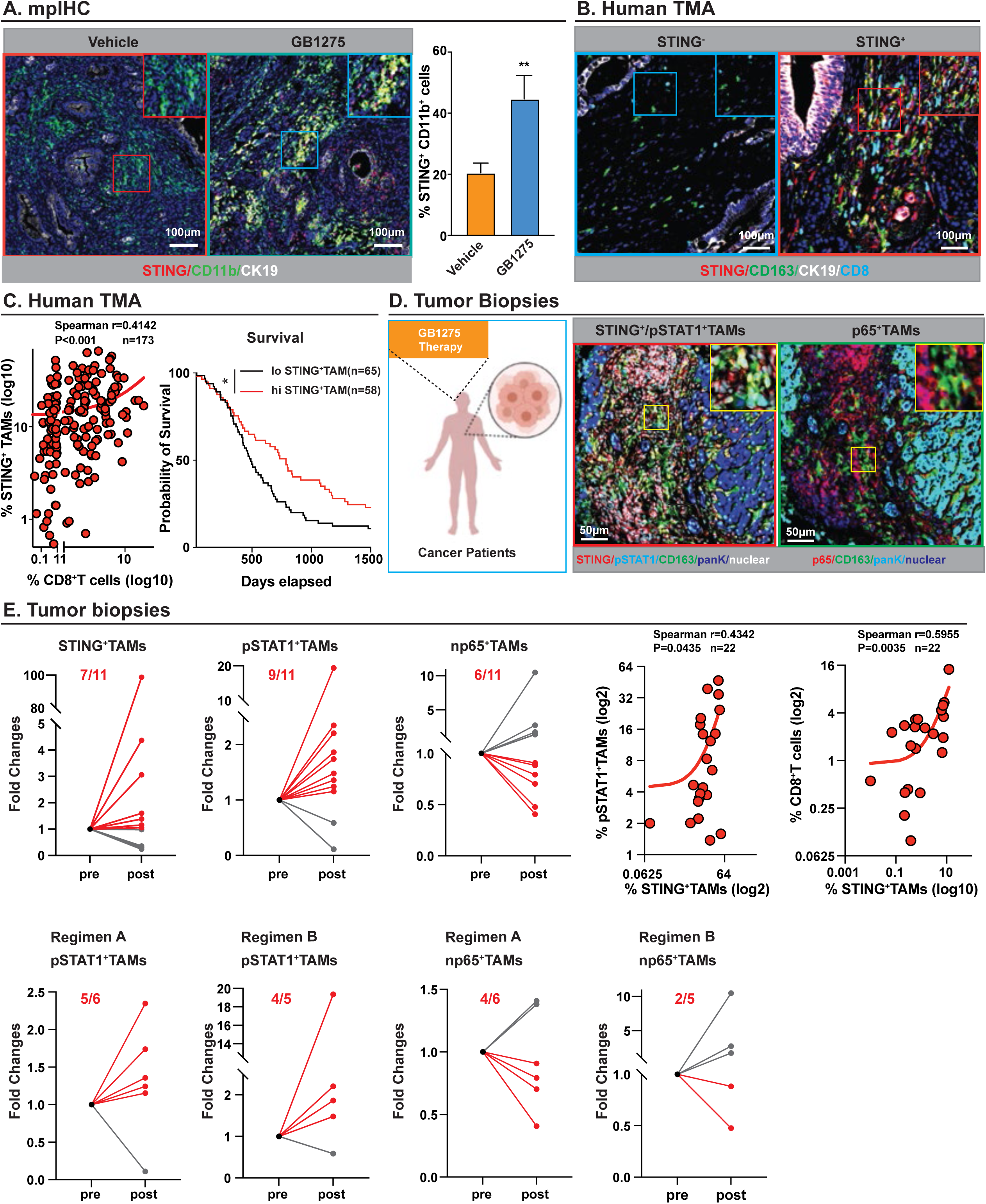
Activated STING/IFN signatures by GB1275 treatment in the mouse model and tumor biopsies. **A**. Representative mpIHC staining for STING, CD11b, and CK19 in tumors from the 14-day vehicle and GB1275 (120mg/kg)-treated KPPC mice. Scale bar, 100 μm. Right, percentages of STING^+^ CD11b^+^ cells from each treatment group (n = 7−9 mice per group). **B**. Representative multiplex immunohistochemistry (mpIHC) staining for STING, CD163, CK19, and CD8α in a human pancreatic cancer tissue microarray. Scale bar, 100 μm. **C**. Scatter plot showing Spearman’s correlation between the percentage of STING^+^ macrophages (out of CD163^+^, CK19^-^ cells) and percentage of CD8^+^ T cells in human pancreatic cancer tissue samples (left). Kaplan-Meier survival curves for patients with high STING^+^TAM infiltration and low STING^+^TAM infiltration from mpIHC analyses (right). **D**. Representative mpIHC staining for STING, pSTAT1, p65, CD163, and pan-keratin (PanK) in 11 paired tumor biopsies from patients. Six pre- and post-pairs were in regimen A (GB1275 monotherapy), five pre- and post-pairs were in regimen B (GB1275+ pembrolizumab). Scale bar, 50 μm. **E**. Relative fold changes of STING^+^ macrophages and pSTAT1^+^ macrophages, nuclear p65+ macrophages in paired pre- and post-groups. Scatter plots showing Spearman’s correlation between the percentage of STING^+^ macrophages (out of macrophages) and pSTAT1^+^ macrophages (out of macrophages) in all 22 tumor biopsies. Scatter plots showing Spearman’s correlation between the percentage of STING^+^ macrophages (out of total cells) and CD8^+^T cells in all 22 tumor biopsies (right). Relative fold changes of pSTAT1^+^ macrophages, nuclear p65^+^ macrophages in paired pre- and post-groups from regimens A and B, respectively (bottom). Graphs show the mean ± SEM; *denotes P < 0.05 using the two-sided *t-*test or log-rank test.

### Biomarker validation in the context of the First-in-Human GB1275 clinical trial

Recently, the first human clinical evaluation of the CD11b agonist, GB1275, in advanced solid tumors was initiated [NTC04060342 and (45)]. The Phase 1 portion of the study included a single agent GB1275 dose escalation and safety evaluation either as monotherapy (Regimen A) or in combination with pembrolizumab (Regimen B). Both treatment regimens included patients with solid tumors refractory to advanced treatments (45). In this ongoing study, GB1275 demonstrated tolerability as monotherapy and combined with pembrolizumab, even at the highest dose level (45). For a subset of patients, pre- and post-treatment tumor tissues were available, and we used these to determine potential biomarkers of pharmacodynamic activity. To determine if GB1275 modulated STAT1/STING and p65 signaling in TAMs in these trial patients, we performed mpIHC for CD11b, CD68, CD163, pSTAT1, STING, p65, and pan-cytokeratin (PanK) on 11 paired, pre-treatment and post-treatment, biopsy tissues (Sup. Fig. 9D, E; Fig. 5D). Compared with pre-treatment tissues, we observed that GB1275 elevated the percentages of pSTAT1^+^ TAMs and STING^+^ TAMs out of total TAMs in the majority of patients across both regimens (9/11 pSTAT1^+^TAMs, 7/11 STING^+^TAMs). The induction of pSTAT1 in post-treatment biopsies appeared independent of exposure to pembrolizumab (5/6 Regimen A, 4/5 Regimen B) (Fig. 5E). Across both regimens, p65 expression in TAMs only decreased in 6 out of 11 patients, and the majority of these patients involved GB1275 as a single agent (4/6 Regimen A, 2/5 Regimen B) (Fig. 5E). This may not be surprising, as activation of STING/STAT1 or other inflammatory pathways *in vivo* during treatment may overcome CD11b agonist’s impact on NFκB (46, 47). As expected, we observed a correlation between TAM STING expression and pSTAT1 activation (Fig. 5E). The percentage of STING^+^TAMs from total cells was positively correlated with CD8^+^ T cell infiltration among all pre- and post-treatment biopsies (Fig. 5E), similar to our data from our PDAC TMA. Taken together, our data revealed that CD11b agonists increased STING/STAT1 activation in TAMs in biopsies from patients with advanced metastatic cancer.

### DNA damaging therapies cooperate with CD11b agonism to amplify STING/IFN signaling in TAMs

The above data demonstrated that CD11b agonists activated STING/IFN signaling in TAMs. However, tumor parameters that might define the magnitude of this activation remain unclear. Notably, *in vitro* GB1275 maximal induction of STING/IFN signaling in macrophages was dependent on TCM (Fig. 4A). Thus, we hypothesized that understanding what factors in TCM augmented STING activation might yield insight into how to further amplify this pathway. To accomplish this aim, we first used a protein concentrator to separate proteins, which were generally larger than 3 kilodaltons (kDa), from small molecules, like metabolites, which are generally smaller than 3 kDa (Fig. 6A). Next, we assayed total TCM, concentrated proteins (> 3 kDa) from the TCM, and metabolites from the TCM (< 3 kDa) for their ability to activate BMDMs in combination with GB1275. Notably, the fraction containing metabolites from the TCM, and not concentrated proteins, enhanced the induction of IFN and CXCL mRNA expressions by GB1275 (Fig 6A). Additionally, these small molecules/metabolites in isolation augmented IFN gene induction by GB1275 better than whole TCM. These data suggested that small molecules or metabolites secreted from tumor cells contributed to GB1275-mediated STING activation. To identify metabolites related with STING signaling, we assessed whole genomic DNA (gDNA) and mitochondrial DNA content in the < 3 kDa TCM fraction. We readily found detectable gDNA and mtDNA in the < 3 kDa fraction of TCM, and the amount of this mtDNA was further increased once TCM was made by tumor cells exposed to cell damaging agents, like gemcitabine (GEM) stimulation (Fig. 6B). To verify the role of gDNA or mtDNA in the TCM, we depleted DNA in the TCM with EtBr and observed that the TCM no longer increased IFN/CXCL by GB1275 treatment (Fig. 6C). These data suggested that even low levels of mtDNA or gDNA from stressed or dying tumor cells augmented STING signaling activation by GB1275. Because GEM-treated PDAC cells released more mtDNA, we determined whether cell damage reagents synergized with GB1275. We found TCM made from PDAC cells exposed to either chemotherapy (GEM) or radiation, while not able to increase IFNβ and CXCL10 gene expression on their own, significantly synergized with GB1275 to induce IFNβ and CXCL10 in macrophages (Fig. 6D). To test whether the above synergism phenomena existed *in vivo*, we treated PDAC-bearing mice with either chemotherapy, GEM plus paclitaxel (GEM/PTX) (Fig. 6E), or radiation therapy (RT) ± GB1275 or in the presence of intratumoral injection of a STING agonist, ADU-S100 (Fig. 6G). Consistent with the clinical experience for PDAC patients, both GEM/PTX and RT only modestly delayed tumor progression; however, when these regimens were combined with GB1275, we observed significant tumor burden reduction and improved overall survival (Fig. 6E, G). We also observed synergy when the STING agonist (AUD-S100) was added to GB1275 and RT (Fig. 6G). In addition, mice in any of these combination groups did not show any body weight loss, and the treatments were well tolerated. Similar to the induction of interferon genes in macrophages observed *in vitro*, analysis of tumor tissues from mice treated with GB1275+chemotherapy showed very significant increases in IFNα, IFNβ, and CXCL9, 10, and 11 mRNAs, when compared with all controls (Fig. 6F). Taken together, our data demonstrated that cytotoxic agents were capable of synergizing with CD11b agonism to amplify STING/IFN signaling activation and tumor control *in vivo*.

**Figure 6.**
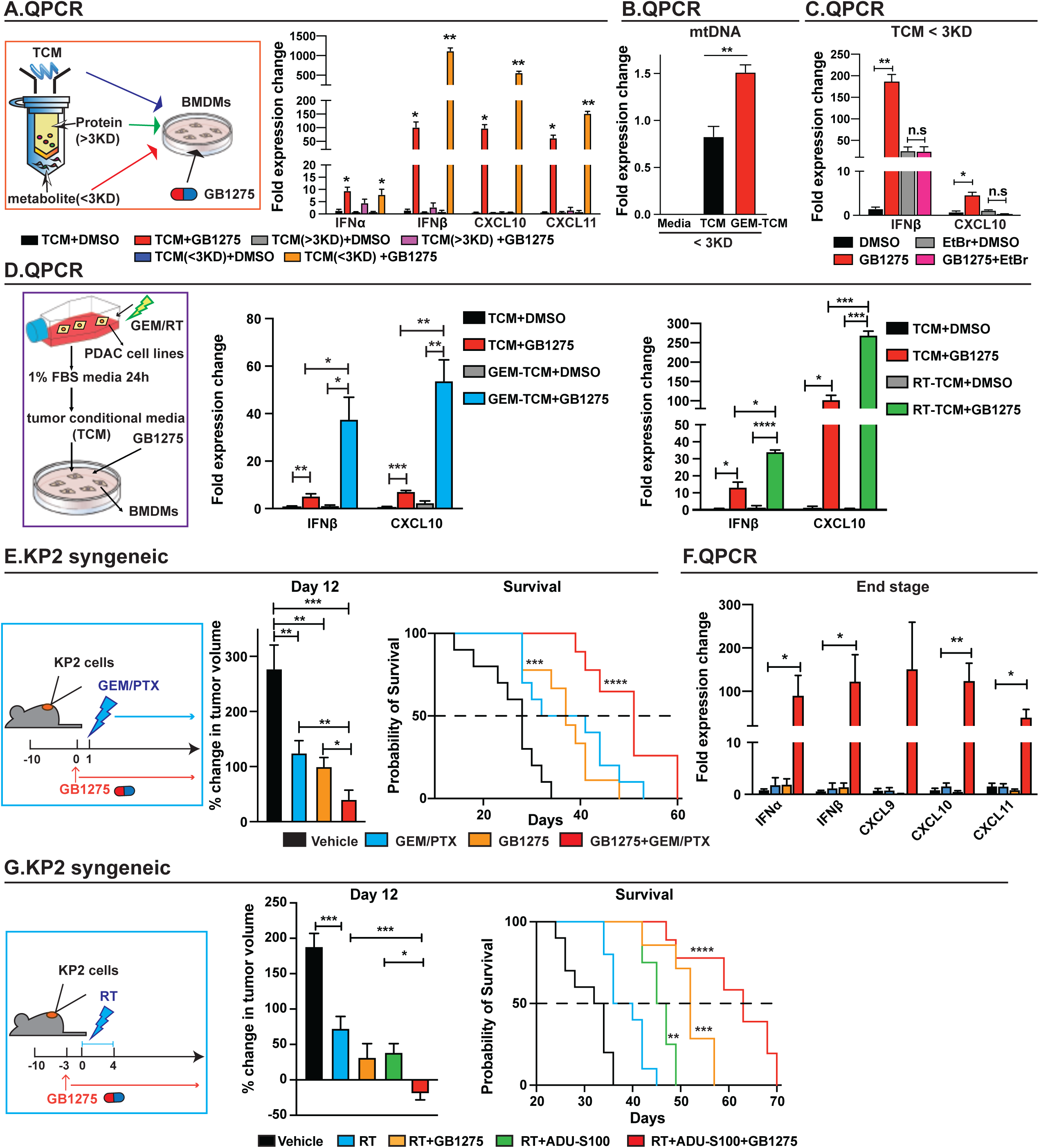
Chemo or radiation, in cooperation with GB1275 amplifies STING/IFN signaling in pancreatic ductal adenocarcinomas. **A**. Concentrated protein (> 3 kDa) and metabolites (< 3 kDa) in tumor conditional media (TCM) were separated by protein concentrator (left). Quantitative PCR mRNA expression analysis of bone marrow-derived macrophages (BMDMs) treated with different fraction of TCM ± vehicle or GB1275 for 7 h. Changes in gene expressions are depicted as the fold change from the vehicle baseline (right). **B**. Cytochrome C released in metabolites (< 3 kDa) from either normal media or TCM (< 3 kDa) made from KP2 cell lines treated with 10 μM gemcitabine or vehicle for 8 h, was detected by quantitative PCR analyses. Changes in gene expression are depicted as the fold change from TCM group baseline. **C**. Quantitative PCR mRNA expression analysis of BMDMs treated with metabolite of TCM (< 3 kDa) in the presence or absence of ethidium bromide + vehicle or GB1275 for 7 h. Changes in gene expressions are depicted as the fold change from the vehicle baseline. **D**. KP2 cells were treated with either 10 μM gemcitabine or 8 Gy radiation. After 8 h, the cells were rinsed with phosphate-buffered saline, and incubated with medium containing 1% fetal bovine serum to make TCM (left). Quantitative PCR mRNA expression analysis of BMDMs treated with the abovementioned TCM ± Vehicle or GB1275 for 7 h. Changes in gene expression are depicted as the fold changes from the vehicle baseline. **E**. Syngeneic tumor growth of KP2 cells in mice treated with vehicle or GB1275 (120mg/kg) ± chemotherapy (50 mg/kg gemcitabine + 10 mg/kg paclitaxel) (left). Mean percent change in tumor volume is shown 12 days after treatment (n = 9–10/group) (middle). (right) Kaplan-Meier survival analysis of mice in these four groups (n = 9–10/group). **F**. Quantitative PCR mRNA expression analysis of tissue from E (n = 6-8 group). Changes in gene expressions are depicted as the fold changes from the vehicle baseline. **G**. Syngeneic tumor growth of KP2 cells in mice treated with vehicle or GB1275 (120mg/kg) ± radiation therapy (6 Gy × 5) in the presence or absence of ADU-S100 (1.25 mg/kg) (left). Mean percent change in tumor volume is shown 12 days after treatment (n = 10/group) (left). (right) Kaplan-Meier survival analyses of mice in these five groups (n =10/group). The graphs show the mean ± SEM; *denotes p < 0.05 using the two-sided *t-*test, analysis of variance, or log-rank test. *In vitro* data are representative of three independent experiments.

### CD11b-agonism enables STING activation to remodel tumor immunity

We next sought to determine the functional consequence of the combination of STING activation in the presence of CD11b-agonist therapy on the tumor immune microenvironment. To accomplish this aim, PDAC-bearing mice were treated with vehicle or GB1275 ± intra-tumoral injection of the STING agonist, ADU-S100. Both GB1275 and STING agonist itself exhibited modest tumor growth suppression, but the combination was superior to either agent alone (Fig. 7A). Both GB1275 and ADU-S100 increased CD8^+^ T cell numbers, but the highest levels of T cell infiltration were seen when STING and GB1275 were combined (Fig. 7B). As expected, that both the STING agonists and CD11b agonists reduced TAM numbers (Fig. 7B), possibly due to macrophage turnover, as previously noted. In KPC GEMMs, we also found a similar increased infiltration of CD8^+^ T cells in PDAC tissues following the combination of GB1275 and STING agonists (Fig. 7C). Together, these data suggested that CD11b agonists and traditional STING activation worked synergistically to regulate T cell infiltration.

**Figure 7.**
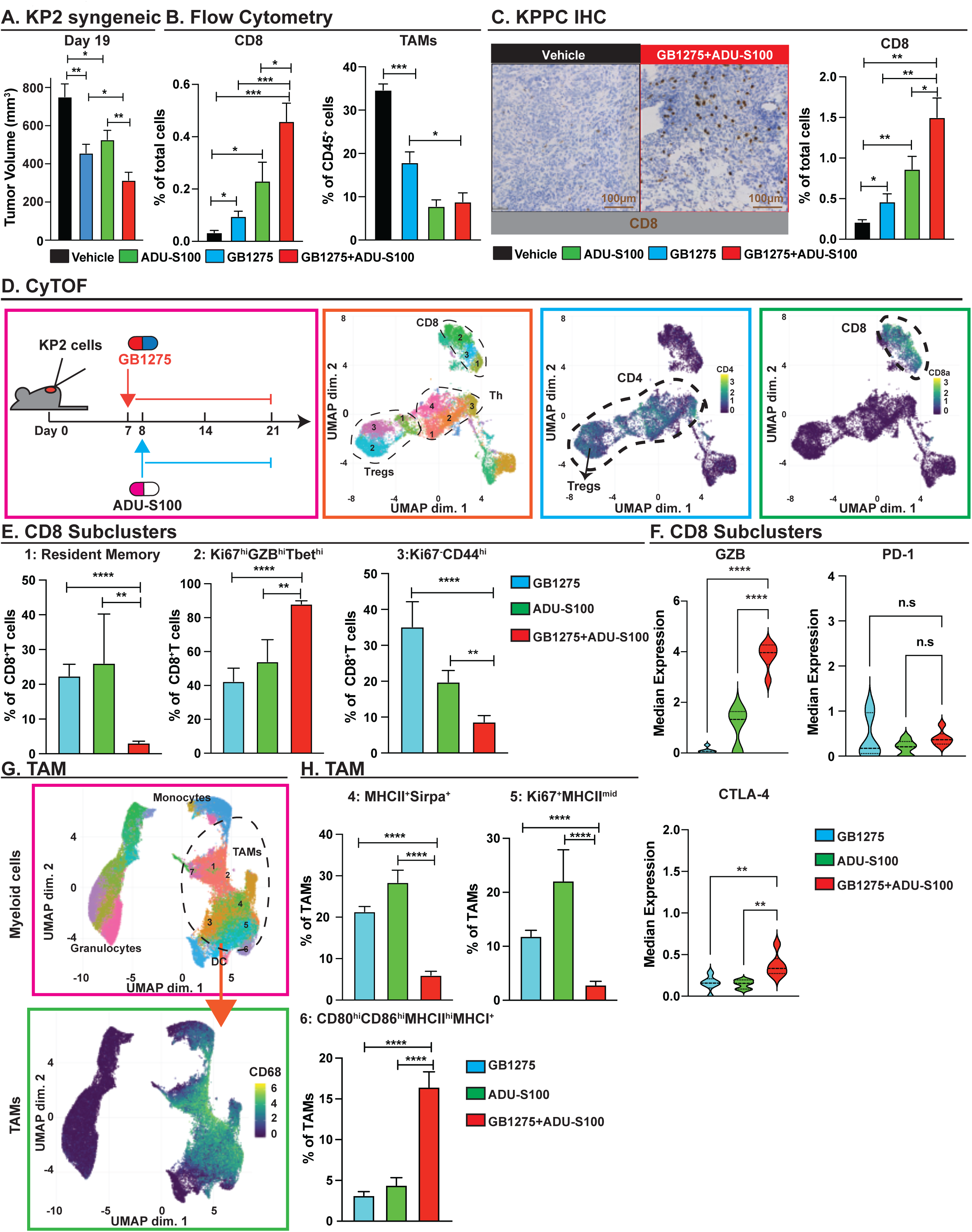
STING agonist synergizing with CD11b agonist remodels the tumor microenvironment. **A**. Tumor burden from the KP2 syngeneic mouse model treated with vehicle or GB1275 (120mg/kg) ± ADU-S100 (1.25 mg/kg) for 19 days (n = 10/group) shown by tumor volume. **B**. Relative frequencies of tumor infiltrating CD8^+^ T cells and macrophages from the above mice (n = 6/group). **C**. Immunohistochemical staining for CD8^+^T cells in tumors from 14-day vehicle and GB1275 (120mg/kg)-treated KPPC mice. Scale bar, 100 μm. Average percentage of CD8^+^ T cells from each treatment group (n = 6−7/group) (right). **D**. Syngeneic model of KP2 treated with GB1275 (120mg/kg), ADU-S100 (1.25 mg/kg), or the combination of GB1275 and ADU-S100 for 14 days (left). CyTOF UMAP plot of tumor infiltrating T cells, including of CD8^+^ T cells, Th, and T^Regs^ from the above tissues (n = 6/group). **E**. Percentages of individual subclusters in CD8^+^T cells from three groups. **F**. Median expressions of GZB, PD-1, and CTLA-4 in CD8^+^ T cells. **G**. CyTOF UMAP plot of tumor infiltrating myeloid cells, including tumor-associated macrophages (TAMs), granulocytes, monocytes, and dendritic cells from the above tissues (n = 6/group). **H**. Percentage of individual subclusters in TAMs from three groups. Graphs show the mean ± SEM; *denotes P < 0.05 using the two-sided *t*-test or analysis of variance.

To more deeply probe the immune changes induced by the combination of CD11b-agonists and STING activation, we used CyTOF (Fig. 7D). As before, UMAP analysis identified several clusters of CD8^+^ T cells, CD4^+^ T effectors, and Tregs (Fig. 7D; Sup. Fig. 10A). Compared to single agent GB1275 or intratumoral ADU-S100, the combination markedly shifted CD8^+^ T cells toward increased proliferative effector Tbet^Hi^/GZB^Hi^/Ki67^Hi^ phenotypes (clusters 2), at the expense of decreases in CD103^+^ resident memory (cluster 1) and non-proliferative Tbet^Mid^/GZB^low^/Ki67^low^ CD8^+^ T cell populations (cluster 3), which were both reduced (Fig. 7E). From expression analyses across all CD8^+^ T cells, ADU-S100 combined with GB1275 significantly increased GZB expression but did not show further upregulation of Ki67 (Fig. 7F; Sup. Fig. 10B). Interestingly, CTLA-4 expression was also enhanced in combination, but not in checkpoints, like PD-1 or LAG3 (Fig. 7F). In CD4^+^Th, compared with both single agents, the combination increased activated CD4^+^ T cells, with low expression of PD-1 (cluster 4). When compared with GB1275, the combination decreased the percentages of proliferative Th cells (cluster 1), and memory CD4^+^ T cells (cluster 2). In contrast with ADU-S100, the combination decreased naïve-like CD4^+^ T cells (cluster 3) (Sup. Fig. 10C). From expression analyses across Th subsets, the checkpoint molecules were not statistically changed by the combination, when compared with single agents (Sup. Fig. 10D). In Tregs, the combination significantly decreased Tregs with high expression of PD-1 (cluster 2) (Sup Fig. 10E), and PD-1 expression in Tregs was downregulated in the combination group, when compared with the GB1275-treated group, while TIM3 expression was elevated. In addition, CTLA-4, TIGIT, and Ki67 expressions in Tregs did not show any differences (Sup Fig. 10F). Together, these data revealed that in addition to T cell numbers, CD11b agonists and STING activation worked synergistically to indirectly enhance CD8^+^ T cells function.

We next analyzed the impact on myeloid phenotypes. UMAP analysis identified several clusters of TAMs, monocytes, and granulocytes (Fig. 7G). Analysis of just the TAMs showed identification of seven subclusters (Sup. Fig. 11A). Of these subclasses, the combination of GB1275 and STING activation decreased Sirpα-high TAMs (cluster 4) and proliferative MHCII-low TAMs (cluster 5), while markedly increasing antigen-presenting TAMs (cluster 6) expressing high levels of CD80, CD86, MHCII, and MHCI (Fig. 7H). These data suggested a shift from phagocytotic and proliferative TAMs toward antigen presentation by the combination, when compared with either single agent alone. Analyzing other myeloid cells, we found that granulocytes had markedly high expression of PD-L1 in combination-treated tumors (Sup. Fig. 11E). Ki67 expression on monocytes was enhanced in the combination group (Sup. Fig. 11D). Interestingly, PD-L1 expression among TAMs (Sup Fig. 11C), granulocytes (Sup Fig. 11E), and conventional dendritic cells (cDCs) (Sup Fig. 11F) was enhanced in the combination group, when compared with both single agents, indicating a possible therapeutic combination with αPD-L1. Together, these data suggested that GB1275 enabled STING activation to more dramatically reprogram tumor immunity to favor T cell responses.

## Discussion

Many cancers are characterized by abundant myeloid cell infiltrates and high numbers of tumor infiltrating myeloid cells often corelate with poor clinical outcomes(48). These myeloid cells include tumor-associated macrophages (TAMs), neutrophils and myeloid-derived suppressor cells (MDSCs) and can drive T cell exclusion and dysfunction in the TME (17,20,49–51). Thus, one promising therapeutic approach is to reprogram myeloid cells to promote T cell-mediated immunity. However, most clinical approaches taken to date have focused on cell depletion. In preclinical models blocking key chemokine receptors (e.g., CCR2 or CXCR1/2) or impairing TAM survival by inhibiting colony stimulating factor-1 receptor (CSF1R) have shown efficacy in slowing progression and improving responses to a variety of agents (15–17). In early stage clinical trials in PDAC patients (e.g. NCT01413022, NCT03496662) some promising early results were observed (52, 53). However, none of these ablative approaches have been enough to translate into late phase clinical success to date. This is likely because these myeloid-ablative strategies are subject to significant compensatory actions by untargeted subsets of monocytes, granulocytes, and/or tissue resident macrophages that may ultimately limit their therapeutic efficacy in humans where efficacy is measured in months and years rather than weeks as in animal models (19, 20). While these data suggest cancer patients might benefit from targeting myeloid cells, current clinical results also highlight the need to improve this strategy. Promising targets for myeloid reprograming include targeting CD40, CD163, CD47/SIRPa, MERTK, TLRs and others (1,48,54). Here-in we investigate CD11b-agonists as a novel myeloid reprograming approach. However, the impact of myeloid cells on tumor immunity is likely divergent across cancers (55) and thus understanding the exact mechanisms of action of each therapeutic strategy is critical to identifying patients who might benefit.

Comparing with melanoma and lung cancer, PDAC might be considered as a “cold” tumor type; and indeed, it exhibits limited response to immunotherapy (56). In these studies, we accessed a cross tumor comparison of TAM phenotypes by scRNAseq and found, that TAMs in PDAC have lower IFN and antigen presentation signatures compared to more immunotherapy-sensitive cancer types like melanoma and lung cancer. In keeping with this assessment, in PDAC and CRC tissues we also observed minimal percentages of STING^+^TAMs and pSTAT1^+^TAMs, suggesting weaker IFN-signaling in TAMs in these cancers. In contrast, IL-1 and tissue remodeling signatures were higher in PDAC in keeping with the stromal desmoplastic response characteristic of this tumor type. Based on these data, we postulate that inducing IFN signaling in myeloid cells may be key to rescuing tumor immunity in some tumor types.

cGAS-STING signaling pathway has been regarded as an important DNA sensing machinery, allowing immune response to infections, inflammation, and cancer(57). However, activated STING plays a complicated role in cancer (58). On one hand, STING activation induces anti-tumor responses via increased interferon secretion and lymphocyte infiltration, which is a promising direction for cancer immunotherapy(59). It was reported that downregulation of STING could be a factor driving resistance to immune effectors in cancer models (60). Conversely, upregulation of STING signaling increases the number of intratumoral CD4^+^ and CD8^+^T cells consistent with upregulated IFN signatures (61). Improved productive CD8^+^T cell cross-priming via Batf3-lineage dendritic cells (DCs) occurs in STING activation condition (62). These experiments validate that activation of the STING pathway and IFN signaling has anti-tumorogenic properties. On the other hand, emerging evidence has indicated the pro-tumoral role of cGAS-STING pathway in some cancer models (63, 64). STING activation-dependent inflammation is supposed to be the major factor driving tumor development (65). Activation of STING results in TANK-binding kinase 1 (TBK1)/ NFkB-dependent inflammatory cytokines production (66, 67), which supports cancer cell growth and confers chemo-resistance (68). Furthermore, disruption of blood vessels has been observed after STING agonist treatment, owing to TNFα secretion (69). The therapeutic anti-tumor function of STING activation could be characterized as enhanced IFN production, while the side effects might be from activated p65-mediated release of inflammatory cytokines. Hence, to overcome it, GB1275 was used in this paper to reprogram TAMs. STING activation was responsible for GB1275-reprogrammed TME, while the compensation of STING activation was ablated, because of p65 degradation by GB1275. Furthermore, combination of GB1275 with a STING agonist exhibited better control of tumor progression, in keeping with the recruitment of more proliferative effectors and antigen-presenting TAMs, while avoiding the NFκB induced inflammatory cytokine response.

DNA released from tumor cells was necessary to fully activate STING signaling primed by CD11b-agonism. Radiation or chemotherapy-mediated cancer cell death was known to cause tumor-derived DNA, including mtDNA and gDNA release (70, 71). Based on that, we employed RT and chemotherapy in combination with GB1275 in PDAC mouse models and found STING/IFN signaling amplified by the combination. Moreover, treatments were well-tolerated, and without any obvious toxicity in the combinations. These data suggest that these combinations might only trigger STING activation in TMEs. This would be an appealing possibility, as direct STING agonists to date have either been limited by toxicity when given systemically or lacked systemic therapeutic effects in metastatic disease when administered intratumorally to a single lesion (72). Additionally, activity of STING agonists has been limited by degradation of cGAMP by extracellular phosphodiesterase and bell-shaped responses, where excessive concentrations of intratumoral STING agonist could lead to T cell apoptosis(73), PD-L1 overexpression on tumor cells and Tregs infiltration, resulting in a negative impact on anti-tumor immunotherapy(74, 75). In addition to this, we are also interested in whether immunotherapy could be synergized with cell damage agents and CD11b-agonists combinations. CyTOF analysis revealed that PD-L1^+^ myeloid cells and CTLA-4 and PD1 expression in CD8^+^ T cells were high CD11b-agonist treated tumors, which also make these combinations intriguing.

Previous studies have demonstrated efficacy of GB1275, a CD11b agonist, in preclinical models (21-23,57), and our data identify that CD11b-agonists reprogram solid tumor immunity through activation of STING-IFN signaling in TAMs. GB1275 entered clinical investigation (NCT04060342) and analysis of STING/pSTAT1 and p65 expression in a limited number of pre- and post-treatment patient tissues are consistent with GB1275 target engagement at the tumor site and suggest the translatability of our mechanistic findings. Specifically, in alignment with the PDAC model data, GB1275 treatment increased presence of STING^+^ and pSTAT1^+^ TAMs in post-treatment biopsies in the majority of samples tested. In parallel, a reduction in p65^+^ TAMs in post-treatment biopsies was observed in a subset of treated patients. Pharmacodynamic response heterogeneity may be attributable to differential TAM phenotypes/functional states across the different indications studied, as highlighted by our scRNAseq analysis of TAMs across different cancers. Alternatively, the observed heterogeneity may be highlighting differences in the wiring of the tumor microenvironment (e.g. presence of mtDNA), as GB1275-mediated STING activation is context dependent. Early clinical studies with GB1275 have shown limited efficacy to date (57). Our data on the GB1275 mechanism of action suggests that the anti-tumor response of CD11b-agonists can be enhanced with DNA damage inducing agents, suggesting that future clinical exploration of GB1275 in combination with chemo- or radiation-therapy may be warranted.

## Materials and Methods

### Human tumor tissue

TMA studies were conducted on surgically resected PDAC specimens from patients diagnosed at the Department of Pathology at Washington University. To assemble TMAs, clearly defined areas of tumor tissues were demarcated, and two biopsies (1.0 mm in diameter) were taken from each donor block. Four µm paraffin sections were used for multiple immunohistochemical analyses. All human tissue studies were approved by the Washington University School of Medicine Ethics Committee. Fully automated image acquisition was performed using a Zeiss Axio Scan Z1 Slide Scanner system with a 20× objective (Carl Zeiss, Jena, Germany) to capture whole slide digital images.

Human tumor biopsy sections from the Phase 1 First-in-Human clinical trial evaluating GB1275 as monotherapy (Regimen A) and in combination with pembrolizumab (Regimen B) in specified advanced tumors (NCT04060342) were provided by Gossamer Bio (45). Following informed consent, core tissue biopsies were obtained pre- and post-GB1275 dose from 11 patients. Post-treatment biopsies were obtained prior to week 8 of treatment. For details on tumor type and treatment see Table in Sup Fig. 9E. Tissues were fixed (10% neutral buffered formalin, 48 hours) and embedded in paraffin immediately after biopsy. Needle core biopsies were fixed for a minimum of 8 hours.

### The genetic mouse PDAC model and other mouse models

KPC (p48-CRE; LSL-Kras^G12D/wt^; p53^flox/flox^) mice were generated in-house, and C57BL/6 breeders were obtained from the Jackson Laboratory (Bar Harbor, ME, USA). KPC mice were backcrossed to C57BL/6 over six generations and validated as C57BL/6 congenic through single nucleotide polymorphism scanning. The CD11b^-/-^ and STING^-/-^ mice (all on the C57BL/6 background) were purchased from Jackson Laboratories. Mice were maintained in the Washington University Laboratory for Animal Care barrier facility, and all studies involving animals were approved by the Washington University School of Medicine Institutional Animal Studies Committee.

### Compounds

GB1275 was provided by Gossamer Bio. GB1275 is an oral CD11b agonist. For animal experiments, GB1275 was administrated at 60 or 120 mg/kg by oral gavage twice a day (BID). GB1275 dosed at 60mg/kg in experiments in Figure 2 and 120mg/kg in all other in vivo experiments. For *in vitro* experiments, 10 μM GB1275 was dissolved in DMSO for BMDMs. STING agonist (ADU-S100) was purchased from MedChemExpress (Monmouth Junction, NJ, USA) and was administered at 1.25 mg/kg by intertumoral injections every 4 days.

### Syngeneic model and preclinical animal cohorts

Syngeneic PDAC tumors were established by surgical implantation, as previously described. Approximately, KP2 (200 K) or KI (100 K) cells in 50 μL Matrigel (BD Biosciences, San Jose, CA, USA) were injected into the pancreas of each mouse. Cohorts of mice were randomized into different treatment groups by gross palpation of tumors in the pancreas. In the transplantable model, 250,000 KP2 cells in 50 μL Matrigel (BD Biosciences) were injected into each mouse’s back/flank. Cohorts of mice were randomized into different treatment groups by tumor volume. Preclinical studies were conducted with 8–15-week-old female mice. Tumor burden was measured by establishing the gross wet weight of the pancreas and comparing it to that of five parallel mice sacrificed at the beginning of treatment. Mice were maintained within the Washington University Laboratory for Animal Care barrier facility. All studies involving animals were approved by the Washington University School of Medicine Institutional Animal Studies Committee.

### Bone marrow transplantation

Six to eight-week-old C57BL/6 mice were exposed to γ-irradiation dosed at 800 Gy (400 Gy×2). Animals were subsequently injected with 5 × 10^6^ BM cells from either WT or STING-null mice. A syngeneic PDAC model was established with these mice after 4 weeks of recovery.

### Mass cytometry

Tumors taken from KI orthotopic models were treated with vehicle or GB1275 for 12 days. Eight tumors from each group were pooled as four samples (two tumors were pooled as one sample). Mouse samples from the KP2 Subq model were treated with GB1275, ADU-S100, or GB1275+ADU-S100. We used six individuals per group. Cells were stained individually for surface and intracellular staining (the antibodies are listed in **Supplementary Table 1**) and fixed overnight as described above. Each sample was then barcoded with a unique combination of palladium metal barcodes using the manufacturer’s instructions (Fluidigm, San Francisco, CA, USA). Following barcoding, the cells were pooled together and incubated overnight in 2% paraformaldehyde containing 40 nM iridium nucleic acid intercalator (Fluidigm) overnight. On the day of acquisition, the barcoded samples were washed and suspended in water containing 10% EQ Calibration Beads (Fluidigm) before acquisition on a CyTOF2 mass cytometer (Fluidigm). Sample barcodes were interpreted using a single cell debarcoder tool. FCS files were then uploaded to Cytobank and manually gated to exclude normalization beads, cell debris, dead cells, and doublets. Classical T cells were classified as CD45^+^, Cisplatin^-^, NK1.1^-^, TCRgd^-^, and TCRb^+^. Myeloid cells were classified as CD45^+^, Cisplatin^-^, and CD90^-^. T cells, and myeloid cells were exported as new FCS files and analyzed using the R CATALYST package^66^ in R (Version 3.8.2: The R Project for Statistical Computing, Vienna, Austria). In brief, FCS files were down-sampled to equivalent cell counts, before clustering with the R implementation of the Phenograph algorithm. Dimensional reduction and visualization were performed using the UMAP algorithm^68^. Finally, differential cluster abundance testing was performed with the R deficit package, utilizing a generalized linear mixed model (GLMM)^69^

### Single cell RNA sequencing data analysis human

Processed count data was downloaded from gene expression omnibus under GSE121636 (76) GSE123814 (77), GSE139555 (78), GSE145370 (79), GSE154826 (80), GSE155698 (81), GSE176078 (82). Ovarian data was downloaded from code repository (83). Expression data was imported into R (v4.1.0) using the Seurat (v4.1.0) R package (84). Cells were filtered for *PTPRC* expression to ensure immune cell. Additional quality control filtering was based on percentage of mitochondrial genes < 10% of counts and removal of cells with feature number greater than 2.5 of the standard deviation of all cells. Myeloid cells were isolated using the scGate (v1.0.0) R package(85) using the “MoMacDC” model. After isolation, manual removal of monocytes utilized feature counts and canonical marker. Dimensional reduction to produce a UMAP (RunUMAP) utilized the standard Seurat workflow, with the addition of data harmonization (RunHarmony) using the harmony (v0.1.0) R package (86) using sequencing run and cancer type as grouping variables Both UMAP calculation and neighbor identification used 15 harmonized dimensions. Single-cell gene enrichment was performed using the UCell implementation (87) in the escape R package (76) across the Hallmark and C2 libraries in the Molecular Signaling Database (88). Statistical testing across all cancer types was performed using Kruskal-Wallis test, with individual comparisons calculated with pairwise Wilcoxon test. Adjusted p-value for significance testing was based on the total number of pairwise Wilcoxon results using the Bonferroni correction for multiple hypothesis testing.

### Single cell RNA sequencing data analysis mouse

PDAC tissues were taken from vehicle-treated, GB1275-treated pancreatic tumors, 14 days post-treatment. Immune cells (CD45^+^) were sorted by an Aria-II cell sorter (BD Biosciences). Each sample was generated from a pool of three mice/treatment group and two total libraries were sequenced.

Sorted cells from each sample were encapsulated into droplets and libraries were prepared using Chromium Single Cell 3’v3 Reagent kits according to the manufacturer’s protocol (10× Genomics, Pleasanton, CA, USA). The generated libraries were sequenced by a NovaSeq 6000 sequencing system (Illumina, San Diego, CA, USA) to an average of 50,000 mean reads per cell. Cellranger mkfastq pipeline (10× Genomics) was used to demultiplex illumine base call files to FASTQ files. Files from the KPC tumors were demultiplexed with > 97% valid barcodes, and > 94% q30 reads. Afterwards, fastq files from each sample were processed with Cellranger counts and aligned to the mm10 reference (Version 3.1.0, 10× Genomics) to generate the feature barcode matrix.

The filtered feature with barcode matrices were loaded into Seurat as objects. For each Seurat object, genes that were expressed in less than three cells and cells that expressed less than 1,000 or more than 6,000 genes, were excluded. Cells with greater than 10% mitochondrial RNA content were also excluded, resulting in 12,960 immune cells in the vehicle and 12,086 in the GB1275-treated group. SCTransform with default parameters was used on each individual sample to normalize and scale the expression matrix against the sequence depths and percentages of mitochondrial genes. Cell cycle scores and the corresponding cell cycle phase for each cell were calculated and assigned after SCTransform based on the expression signatures for S and G2/M genes (CellCycleScoring). The differences between the S phase score and G2/M score were regressed-out by SCTransform on individual samples. Variable features were calculated for each sample independently and ranked, based on the number of samples they were independently identified (SelectIntegrationFeatures). The top 3,000 shared variable features were used for multi-set canonical correlation analysis to reduce dimensions and identify projection vectors that defined shared biological states among samples and maximized overall correlations across datasets. Mutual nearest neighbors (MNNS; pairs of cells, with one from each dataset) were calculated and identified as “anchors” (FindIntegrationAnchors). Multiple datasets were then integrated based on these calculated “anchors” and guided order trees with default parameters (IntegrateData). Multiple datasets were then integrated based on using Harmony Integration (RunHarmony). Principle component analysis (PCA) was performed on the 3,000 variable genes calculated earlier (function RunPCA). A UMAP dimensional reduction was performed on the scaled matrix using the first 30 PCA components to obtain a two-dimensional representation of cell states. Then, these defined 30 dimensionalities were used to refine the edge weights between any two cells based on Jaccard similarity (FindNeighbors), and were used to cluster cells through FindClusters functions, which implemented shared nearest neighbor modularity optimization. To characterize clusters, the FindAllMarkers function with log-fold threshold = 0.25 and minimum 0.25-fold difference and MAST test results were used to identify signatures alone with each cluster. Then, the TAMs were selected, and the top 3,000 variable features were recalculated to re-cluster. DEGs between the two groups were calculated for each dataset with min.pct of 0.1 and logfc. threshold of 0.01 and MAST test (FindMarkers). Then, the DEG lists from each dataset were filtered with a value of P < 0.05 and ranked based on fold change. These ranked gene sets were fed into GSEA to test for GO terms, KEGG pathways, Reactome Database, and the Molecular Signatures Database gene sets with false discover rate (FDR) < 0.05.

### Mouse tissue isolation and flow cytometry

Mice were euthanized by intracardiac perfusion using 20 mL of PBS-heparin. Tumor tissues were manually minced and digested in 25 mL of Dulbecco’s Modified Eagle Medium (DMEM) containing 2 mg/mL of collagenase A (Roche) and DNase I (Sigma-Aldrich, St. Louis, MO, USA) for 30 minutes at 37°C. Digestion was quenched in 5 mL of fetal bovine serum and filtered through 40 μm nylon mesh. Single cell suspensions were subsequently labeled with fluorophore-conjugated anti-mouse antibodies at recommended dilutions following the manufacturers’ recommendations (**Supplementary Table 2**). Data were acquired on a X-20 (BD Biosciences) and analyzed using FlowJo software (Tree Star).

### Immunohistochemistry and mpIHC

Tissues were fixed in formalin for 24 h and embedded in paraffin according to standard protocols. Immunostaining was performed on the Leica Bond III RXm autostainer using Leica Bond ancillary reagents and a Refine Polymer DAB detection system. Primary antibodies are listed in (**Supplementary Table 3**). For mpIHC of mouse PDACs, embedded tissues were sectioned into 6 um sections and loaded into BOND RXm (Leica Biosystems) for a series of stainings, including antibodies to STING, P65, CD11b, and CK19. Human TMA slides were stained with antibodies to CD8, STING, CD163, CD11b, and CK19. Tumor biopsy slides were stained with antibodies to CD8, STING, pSTAT1, CD163, CD11b, and PanK. Based on antibody host species, default manufacturer protocols were used (IntenseR and Polymer Refine), containing antigen-retrieval with citrate buffer, goat serum and peroxide block, primary antibody incubation, post-primary incubation, and chromogenically visualized with an AEC substrate (Abcam, Cambridge, UK). Between each two cycles of staining, the slides were manually stained for hemoxylin and eosin, then scanned by Axio Scan.Z1 (Zeiss, Jena, Germany). The slides were then destained by a gradient of ethanol plus a 2% hydrochloride wash and blocked with extra avidin/biotin (Vector Laboratories, Burlingame, CA, USA) and a Fab fragment block (Jackson Laboratory, Bar Harbor, ME, USA).

Images of the same specimen, but using different stains, were cropped into multiple segments by Zen software (Zeiss). Each segment was then deconvoluted (Deconvolution, Version 1.0.4; Indica Labs) for individual stains and fused using HALO software (Zeiss) with the default manufacturer’s settings. Markers of interest were pseudo-colored and quantified using High Plex FL software, version 4.0.3 (Indica Laboratories, Albuquerque, NM, USA).

### Macrophage depletion

To deplete tissue resident macrophages, 6–8-week-old FVBN mice were treated with three doses of CSF1 neutralizing antibody (clone 5A1; BioXCell, Lebanon, NH, USA) (1 mg, 0.5 mg, and 0.5 mg on days 3, 10, and 17; Figure 1F) and three doses of clodronate-containing liposomes (Liposoma; 200 μL/each on days 4, 11, and 18). Control mice were treated with same doses/volume of IgG (clone HRPN, BioXCell) and control liposomes (or PBS as indicated, Liposoma). On day 0, mice were implanted orthotopically with 100,000 KI cells and then treated with GB1275 as previous described.

### Radiation therapy (RT)

Ten days post-tumor implantation, cohorts of mice were randomized into different treatment groups using either gross tumor diameters or tumor volumes [length × (width^2^)/2]. Mice were given daily fractionated doses of RT (6 Gy × 5) using the Small Animal Radiation Research Platform (SARRP200; XStrahl Life Sciences, Suwanee, GA, USA). Mice were placed on the irradiation platform one at a time and fitted with a nose cone for isoflurane anesthesia. Cone beam computed tomography imaging was performed for each individual mouse to pinpoint the pancreas tumors, and the images were imported into Muriplan and used to select an isocenter. The tumor was then irradiated to 6 Gy using anterior-posterior-opposed beams using the 10 mm × 10 mm collimator at a dose rate of 3.9 Gy/min.

For *in vitro* radiation experiments, a RS2000 160 kV X-ray irradiator using a 0.3 mm copper filter (Rad Source Technologies, Buford, GA, USA) was used.

### Statistical analyses

All statistical analyses were performed using GraphPad Prism software, with input from a Biostatistics Core expert at Washington University. All data are representative of at least two independent experiments, unless specifically noted. Sample size was pre-calculated to satisfy power requirements (with > 85% confidence) in most experiments and is specified in the figure legends wherever applicable. To accomplish randomization for orthotopic or syngeneic tumor experiments, animals were sorted by a blinded investigator, with tumor sizes in ascending order and then the groups were assigned in descending order. Each group was checked post-hoc to verify that there was no statistical difference in average starting tumor size. Data are shown as the mean ± SEM, unless otherwise noted. Statistical tests such as unpaired parametric Student’s *t*-test, analysis of variance (Bonferroni multiple comparison), or the unpaired nonparametric Mann-Whitney U test was used based on the normality of data. For survival analyses, the log-rank (Mantel-Cox) test was used. For proximity analyses, the nonparametric Kolmogorov-Smirnov test was used to distinguish differences in frequency distributions. A value of p < 0.05 was considered statistically significant for all studies; n.s. denotes not significant.

## Data availability

The scRNA sequencing data from PDAC lesions was found at the Gene Expression Omnibus Repository (GEO) accession number #####. All software packages used are publicly available through commercial vendors.

## Authors’ Contributions

DGD and XL conceived and designed the project. Wet lab experiments and analysis were performed by XL, BLK, JMB, CZ, VL, CM and KL. Dry lab analysis was performed by XL, GDH, JMB, VK, NB and CZ. Technician experience, critical reagents and experimental design input was provided by CM, JKS and VG. Clinical trial samples and information was provided by HP, RO, AG, JMB and LC. The manuscript was written or edited by XL, DD, RO, AG LC and VG.

## Acknowledgments

DGD was supported by NCI R01CA273190 R01CA177670, R01CA203890, P30CA09184215, P50CA196510, and the BJC Cancer Frontier Fund. The author would like to thank Gossamer Bio for providing reagents, clinical samples and advice. The author would also like to thank the CHiiPs Immunomonitoring Laboratory for CyTOF experiments, the Flow Cytometry & Fluorescence Activated Cell Sorting Core for sorting and flow cytometry experiments, and the Genome Technology Access Center for scRNAseq and microarray experiments.

