## supplementary figures and methods for "Context-dependent triggering of STING-interferon signaling by CD11b agonists supports anti-tumor immunity in mouse models and human cancer patients"

#### Cell lines and Compounds

KP2/KP1 cells were derived from a KPC tumor obtained in house. Kras-INK (KI) cells were obtained from Dr. Douglas Hanahan's laboratory. Panc-1, Capan-1, CF-PAC, H-PAC cells were obtained from Dr. Kian H. Lim's laboratory. All cell lines tested negative for MAP and mycoplasma.

Gemcitabine hydrochloride was purchased from Ark Pharm, Inc. and was administered at 50mg/kg by Intravenous injection (i.v) every five days. For in vitro experiment, Gemcitabine was treated at 10μM dissolved in PBS for BMDMs. PACLITAXEL INJECTION was purchased from FRESENIUS KABI and was administered at 10mg/kg by Intravenous injection (i.v) every five days. Autophagy inhibitor, 3-MA was purchased from Sigma-Aldrich. N-Acetyl-L-cysteine was purchased from Sigma-Aldrich. FAK inhibitor (FAKi) was provided by by Verastem Inc.

#### Isolation of Bone marrow- derived macrophages

BM-MACs were isolated following the protocol described previously (27). Marrow cells were isolated by flushing femurs and tibias from C57BL/6J mice and cultured in DMEM/F12 medium (Lonza) containing 10% FBS, penicillin/streptomycin (Gibco) and 20 ng/mL macrophage colony–stimulating factor (M-CSF, PeproTech). After 7 days in culture, adherent macrophages were harvested and seed on 10μg/mL Fibronectin (Sigma Aldrich)- coated plate for different experiments. Macrophages were pretreated with GB1275 and then co-cultured with TCM for 7 hours prior to RNA isolation.

#### RNA isolation and real time PCR

Total RNA was extracted from tissue or cells, using an E.Z.N.A. Total RNA Kit (OMEGA). Complementary DNAs (cDNAs) were synthesized using qScript cDNA SuperMix (QuantaBio). Quantitative real-time PCR Taqman primer probe sets (Applied Biosystems) were used

(**Supplementary table 5**), and the relative gene expression was determined on an ABI 7900HT quantitative PCR machine (ABI Biosystems) using Taqman Gene Expression Master Mix (Applied Biosystems). The comparative threshold cycle method was used to calculate fold changes in gene expression, which were normalized to the expression of glyceraldehyde-3-phosphate dehydrogenase (GAPDH) TATA-box binding protein (TBP) and/or hypoxanthine phosphoribosyl transferase (HPRT) as reference genes.

**Reverse Phase Protein Array (RPPA)**

Cell extracts were lysed using (RIPA) lysis buffer [25 mM Tris-HCl pH 7.5, 150 mM NaCl, 1% NP-40, 0.5% DOC, 0.1% SDS] supplemented with protease and phosphatase inhibitors (Roche). Samples were then submitted to the MD Anderson Cancer Center for the RPPA assay (RPPA CORE 11192019_169).

**Western immunoblot and Immunoprecipitation (IP)**

Cell lysates were harvested using radioimmunoprecipitation assay (RIPA) lysis buffer [25 mM Tris-HCl pH 7.5, 150 mM NaCl, 1% NP-40, 0.5% DOC, 0.1% SDS] supplemented with protease and phosphatase inhibitors (Roche). Cell lysates were resolved in Tris-glycine sodium dodecyl sulfate/polyacrylamide gel electrophoresis (SDS/PAGE) gels and transferred to polyvinylidene difluoride (PVDF) membranes (Invitrogen). After blocking in 1X TBST buffer with 5% w/v BSA, membranes were probed with primary antibodies against pp65, tp65, pIκB, tIκB, STING, pIRF3, tIRF3, pSTAT1, pP38, pMK2 and β-actin overnight at 4°C (**Supplementary table 4**). Membranes were washed thrice in 1X TBST and probed with HRP-conjugated secondary antibody for 1 hour at RT. Membranes were developed with Pierce ECL Western Blotting Substrates and detected using a ChemiDoc XRS+ system (Bio-Rad). For IP, cell lysates were incubated with p65 antibody overnight at 4 °C and then following protein A agarose (Cell Signaling

Technology) incubation for 3 hours. The beads were washed 5 times with lysis buffer and then used for western immunoblot.

**Isolate mtDNA release**

mtDNA was isolated following the protocol described previously(89). Briefly, after different treatment, cells were lysed by 1% NP40 for 15 min on ice. lysates were spined at 13,000 rpm for 15 min at 4 °C. Supernatant was transferred to isolated mtDNA. NucleoSpin Tissue Kit (Takara Bio USA, Inc) was adopted to purity mtDNA from the cytosolic fraction according to the manufacture's instruction. After extracting DNA from the cytosolic fraction, Q-PCR was employed to measure cytosolic mitochondrial DNA. The comparative threshold cycle method was used to calculate fold changes in gene expression, which were normalized to the expression of as 18S rRNA reference genes.

**Measure ROS production**

Briefly, after different treatment, cells were harvested and incubated with 10μM 2′-7′-dichlorodihydrofluorescein diacetate (DCFH-DA) (Sigma- Aldrich) attenuated with serum-free medium for 20 min at 37°C in the dark. After incubation, samples were washed twice with cold PBS. Data were acquired on X-20 (BD Biosciences) and analyzed using FlowJo software (Tree Star).

**Cell proliferation assay**

Cell proliferation assay was performed by using CellTiter96 Non-Radioactive Cell Proliferation Assay (Promega) according to manufacturers' instructions. Briefly, 20,000 BMDMs/ well were seeded into 96-well plates and incubated overnight. After different treatment, Dye

Solution was added to live cultures for 4 h at 37 °C. Absorbance was measured at 570 nm on Multiskan GO plate reader (Thermo Fisher Scientific).

**In vitro Co-culture and siRNA treatment**

Small interfering RNAs (siRNAs) targeting mouse RelA, and STING were purchased from Integrated DNA Technologies (IDT). Sequences are as listed **Supplementary table 6**. siRNA transfections for primary BMDMs were performed using the Mouse Macrophage Nucleofector™ Kit (Lonza) and Nucleofector™ 2b Device (Lonza) with pre-written program Y-001 for BMDMs following the manufacturer's instructions. RNAs and proteins from transfected primary cells were harvest 24 hours after the transfections.

**Isolation of human CD14$^+$ monocytes-derived macrophages**

Leukoreduction chambers from normal donor were obtained from the BJH Pheresis Center (Washington University). Human peripheral blood mononuclear cells (PBMCs) were isolated using Dr. Fehniger's lab protocol (Washington University). Briefly, chamber eluate was mixed well with PBS containing 1 unit/mL heparin, centrifuged with brakes-off and 'buffy layer' was isolated. Tube was centrifuged at 1,800 rpm for 10 min and pellets were incubated in 1X RBC lysis buffer. After subsequent centrifugation at 1,300 rpm for 4 min, PBMC pellets were resuspended in RPMI containing 10% human AB serum (Sigma- Aldrich). Human CD14$^+$ monocytes were sorted from PBMCs using EasySep Human CD14 Selection Kit (Stemcell Technologies Inc.) per manufacturer's instructions. CD14$^+$ monocyte isolation purity > 90% was confirmed by flow cytometry. CD14$^+$ monocytes were cultured in RPMI medium containing 10% FBS, penicillin/streptomycin (Gibco) and 50 ng/mL macrophage colony–stimulating factor (M-CSF, PeproTech) on 10μg/mL Fibronectin (Sigma Aldrich)- coated plates. After 7 days in culture, adherent macrophages were used for different experiments.

**Separation of TCM**

10 mL TCM was filled into the top tube of protein concentrators (3K MWCO, Thermo Scientific™), and spined at 4600g for 1h at 4 °C, until 80%-90% percentage of solution from top tube went in the bottom tube. Concentrated proteins (>3KD) were in the top tube, and the metabolites (<3KD) was in the bottom tube. The different parts of TCM was used to treat BMDMs.

### Supplementary Figure 1

## A.

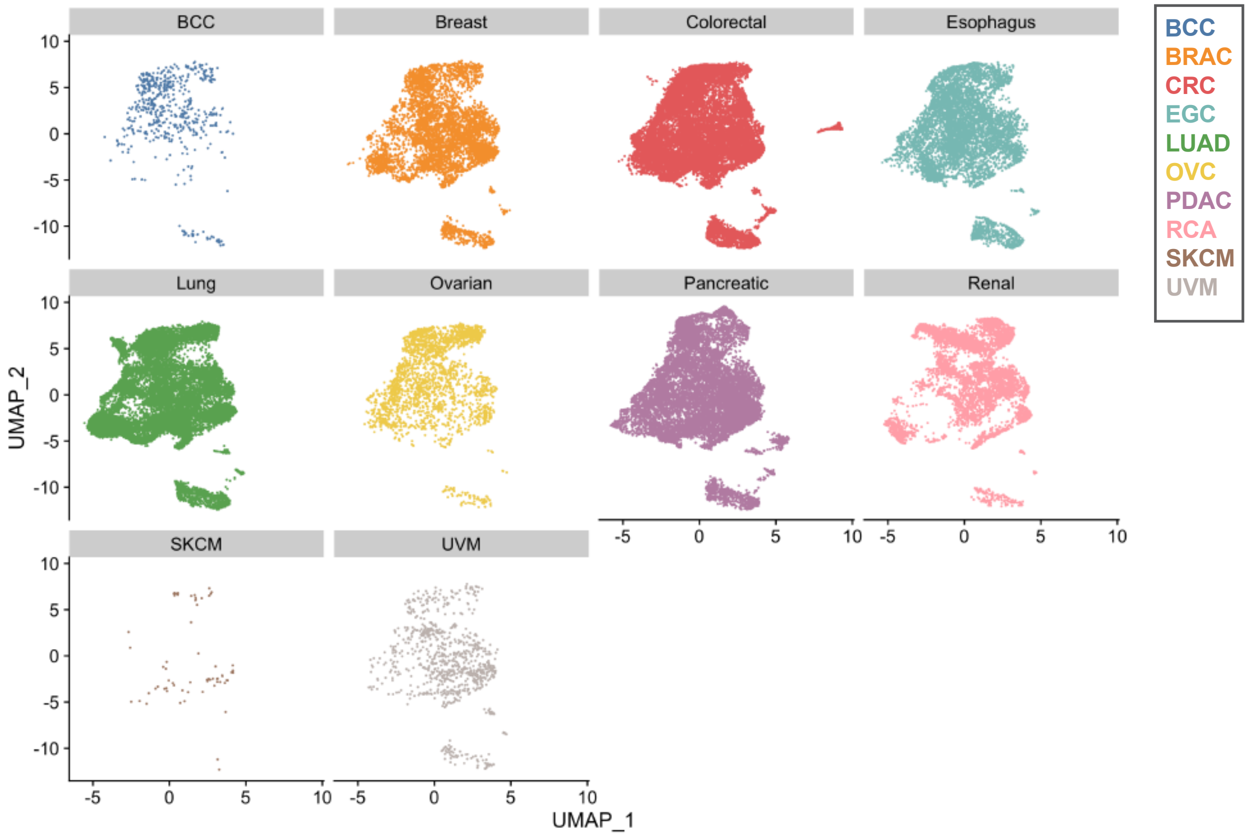


#### B. GSEA Heatmap PID

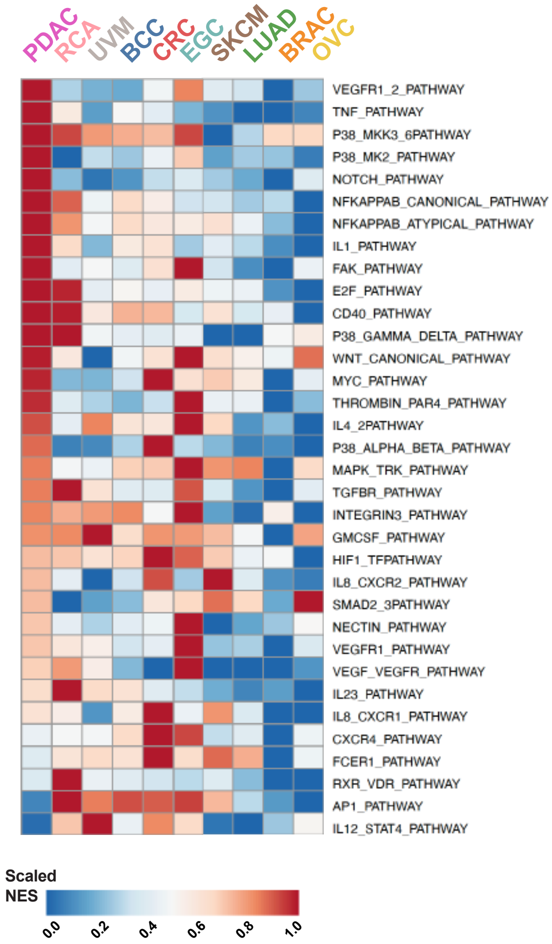


#### C. GSEA Heatmap BIOCARTA

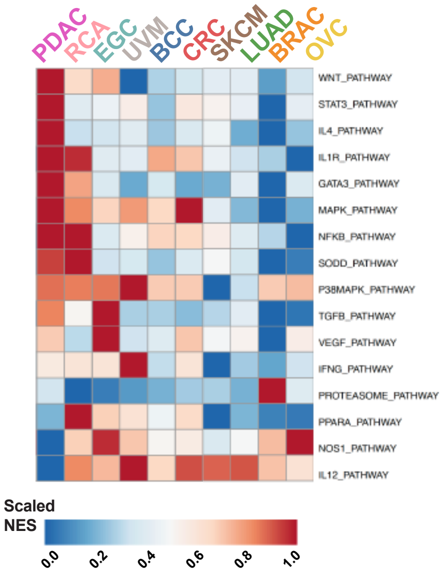

### Supplementary Figure 2

#### A. CD11B agonists

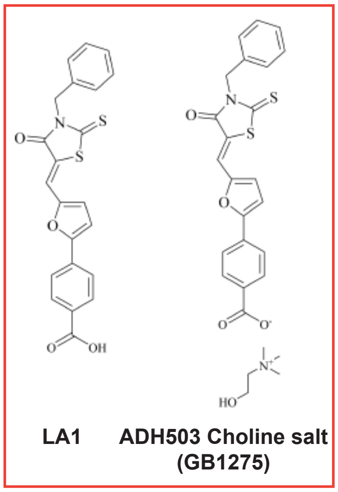


#### B. KP- Orthotopic model

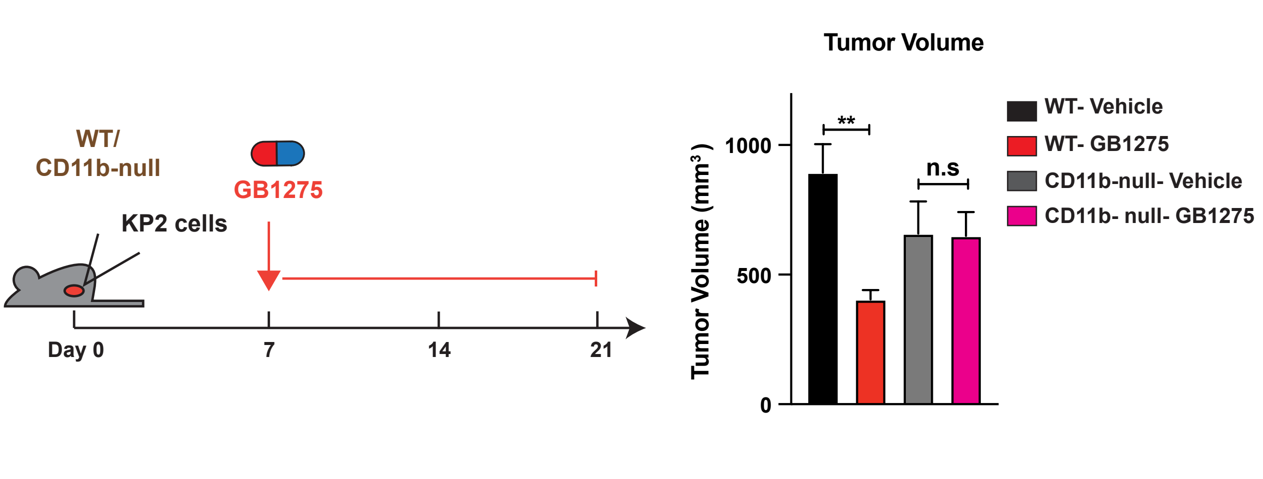


#### C. Flow Cytometry: Myeloid cells

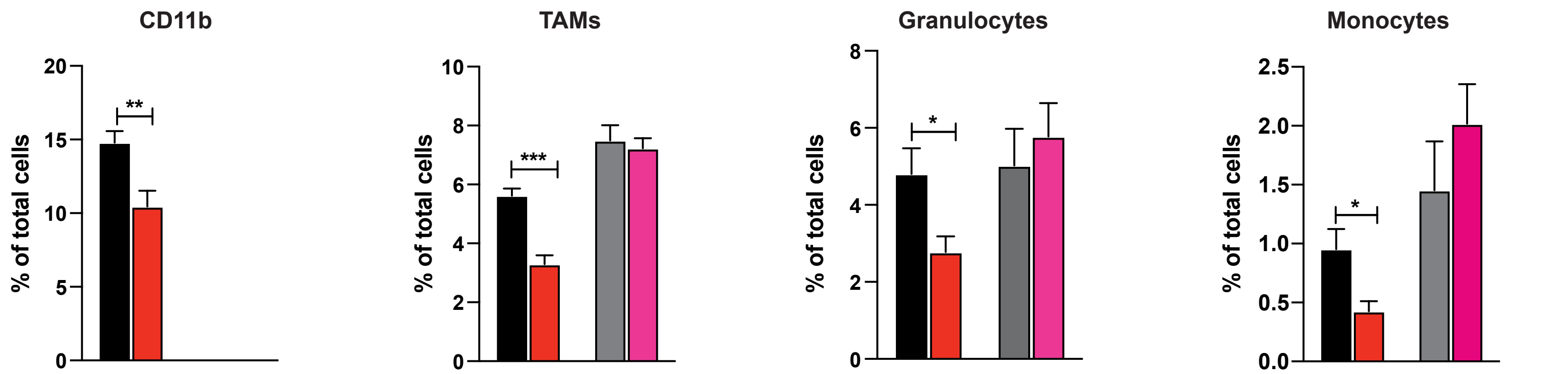


#### D. Flow Cytometry: T cells

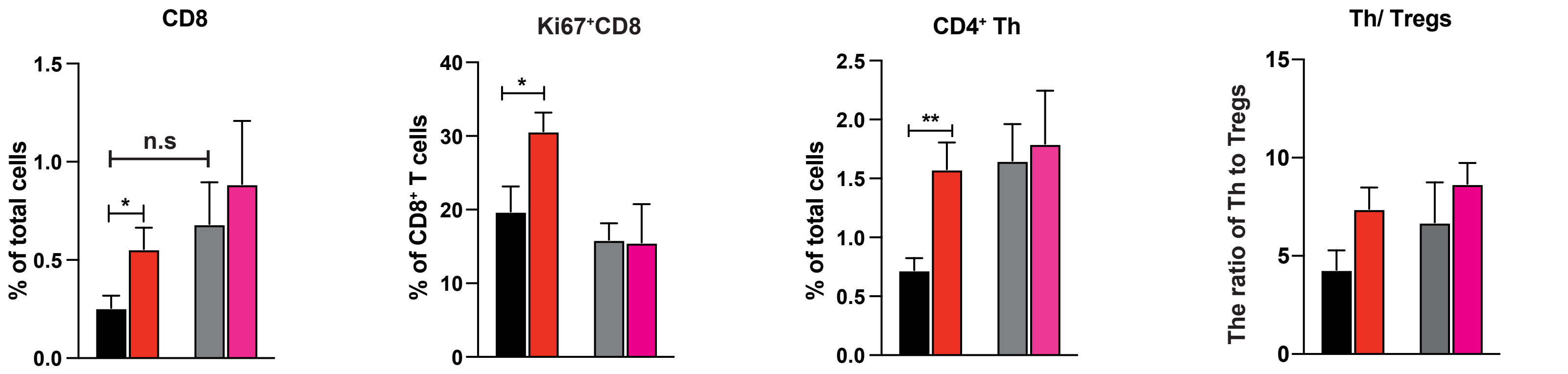

### Supplementary Figure 3

#### A. Gate strategy (Myeloid cells)

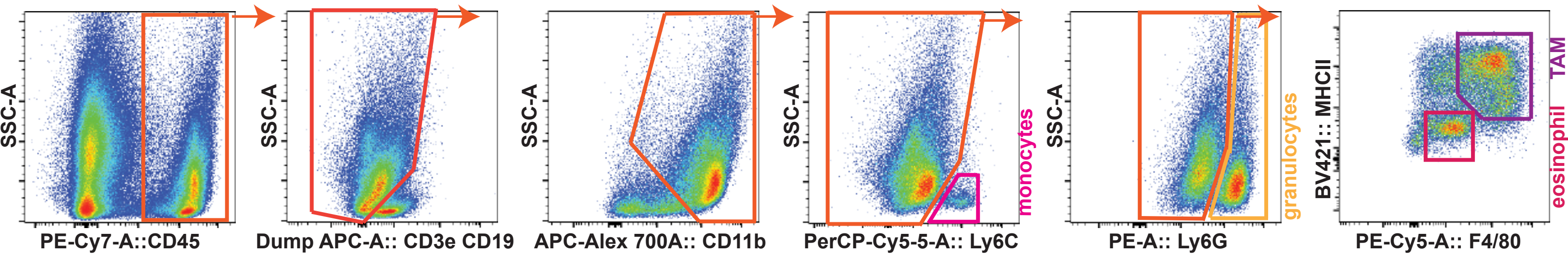


#### B. Gate strategy (T cells)

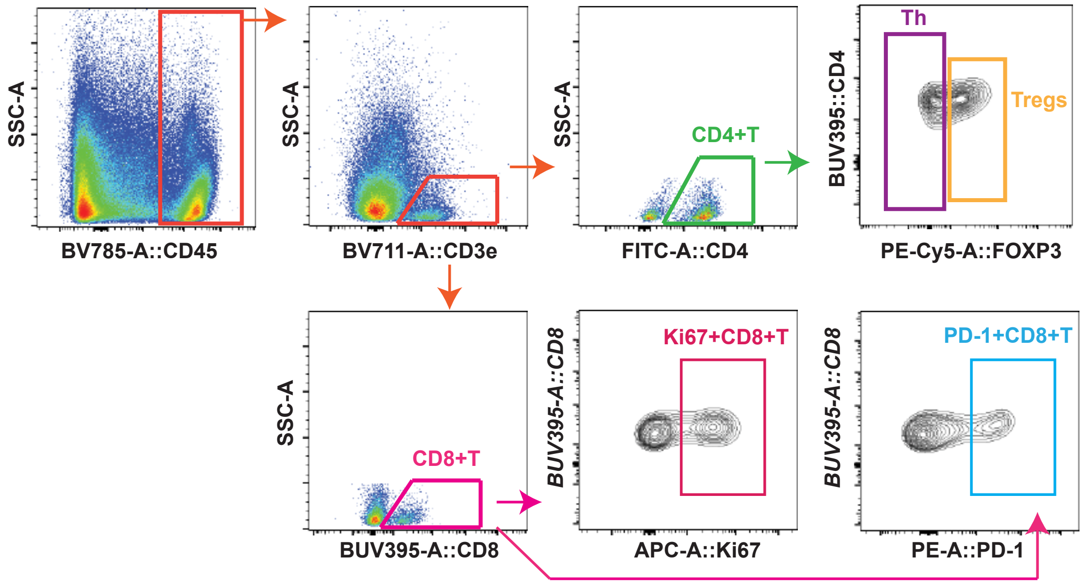

#### Supplementary Figure 4

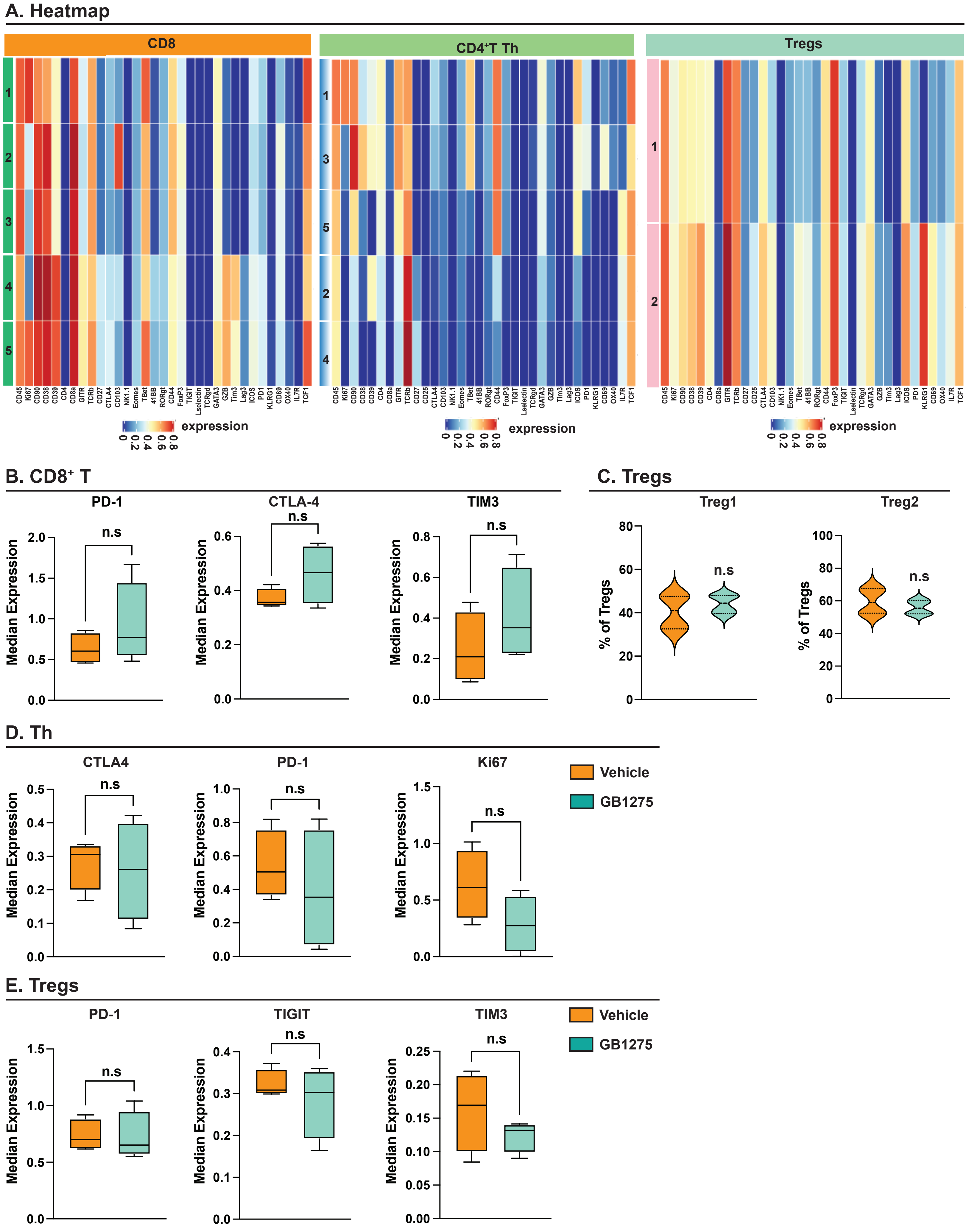

#### Supplementary Figure 5

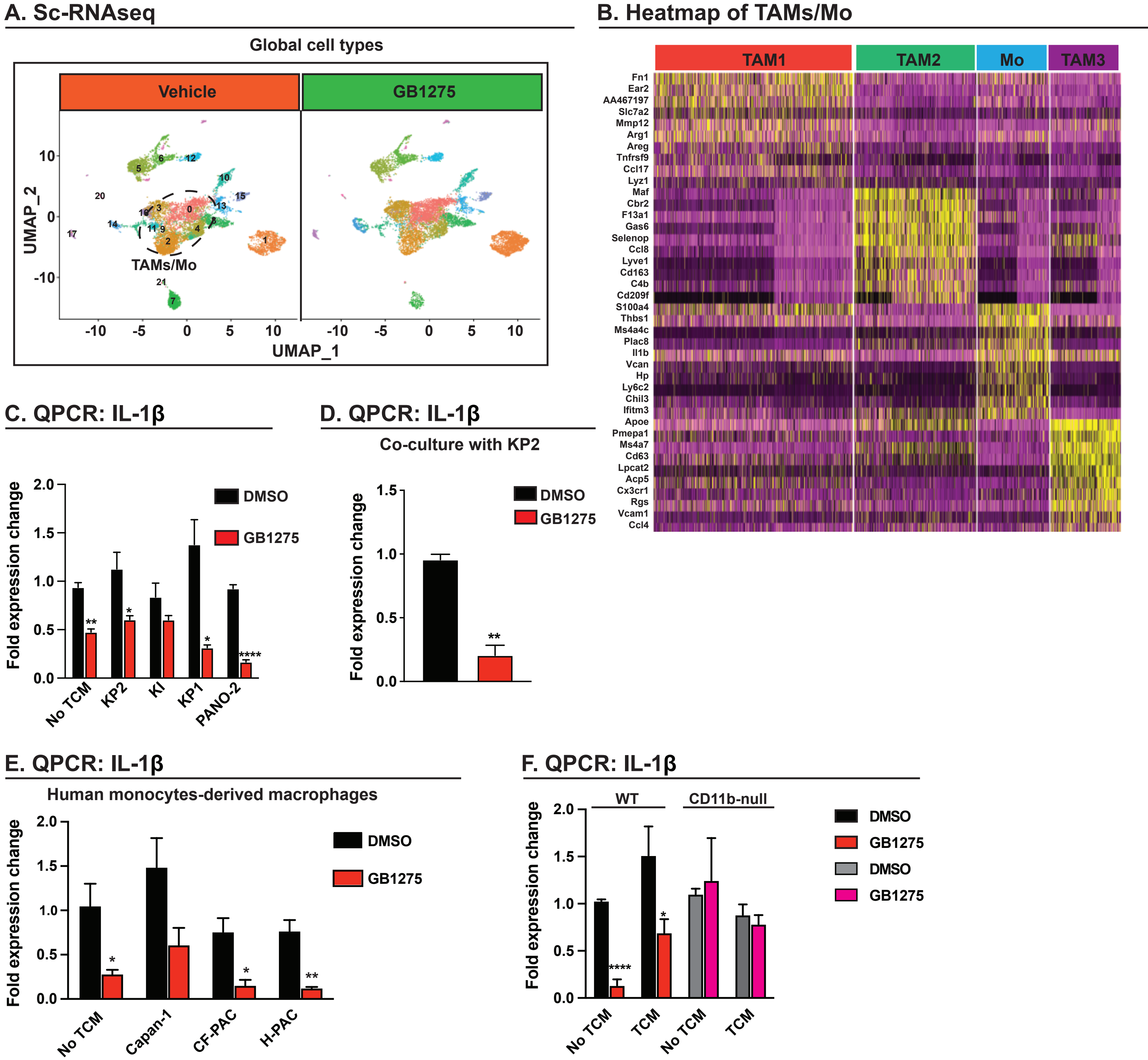

#### Supplementary Figure 6

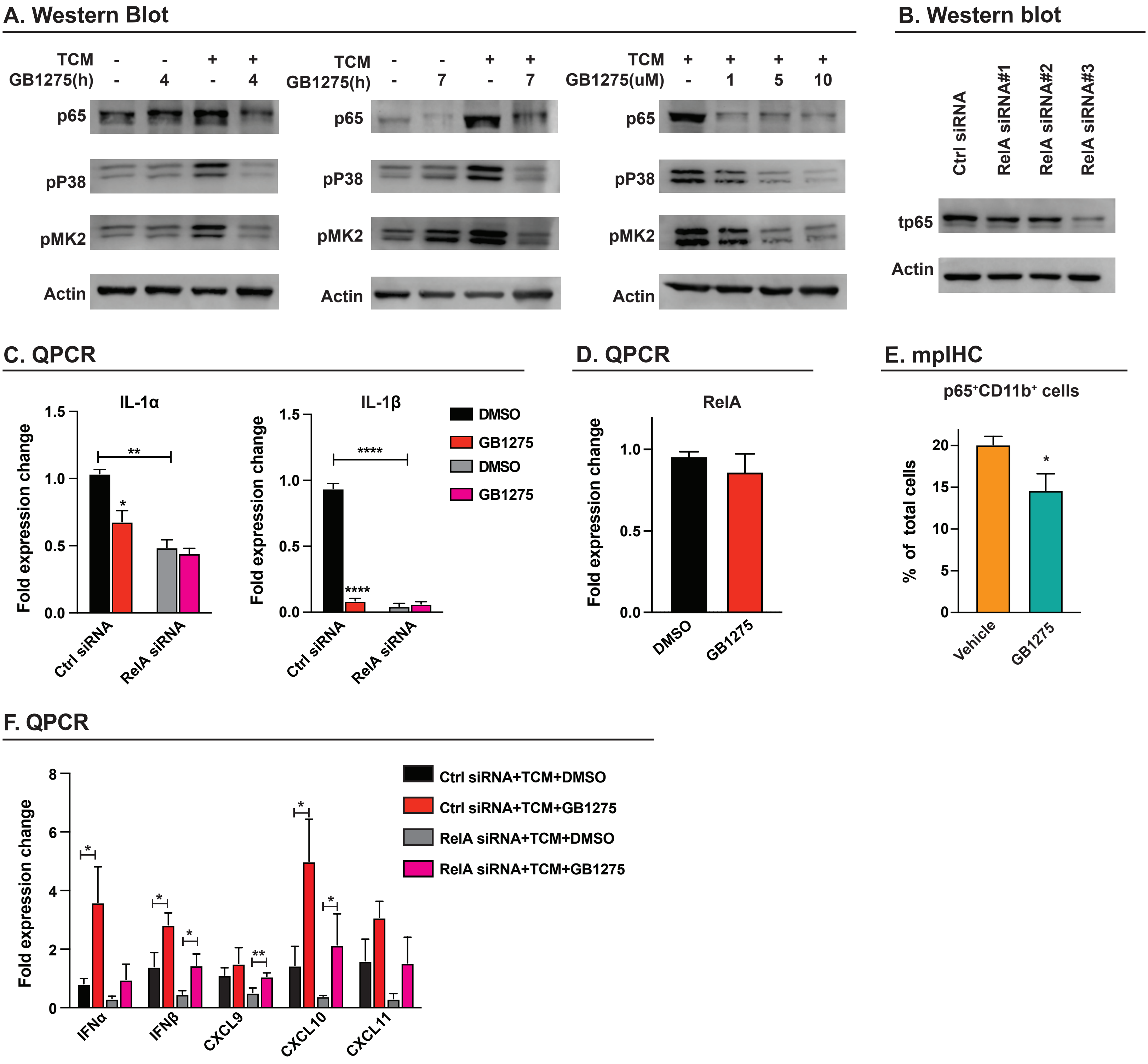

#### Supplementary Figure 7.

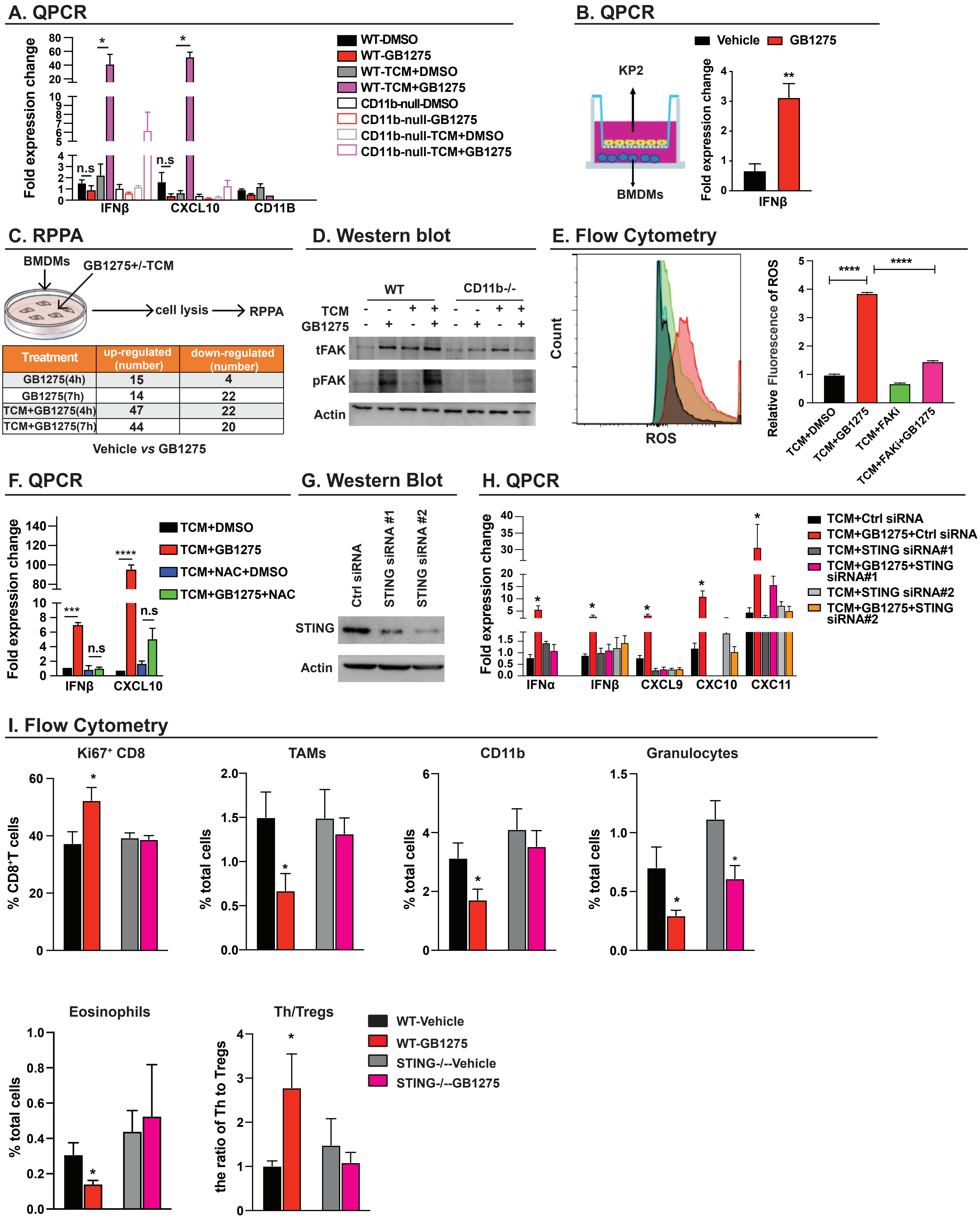


| Treatment | up-regulated (number) | down-regulated (number) |
|---|---|---|
| GB1275(4h) | 15 | 4 |
| GB1275(7h) | 14 | 22 |
| TCM+GB1275(4h) | 47 | 22 |
| TCM+GB1275(7h) | 44 | 20 |

#### Supplementary Figure 8.

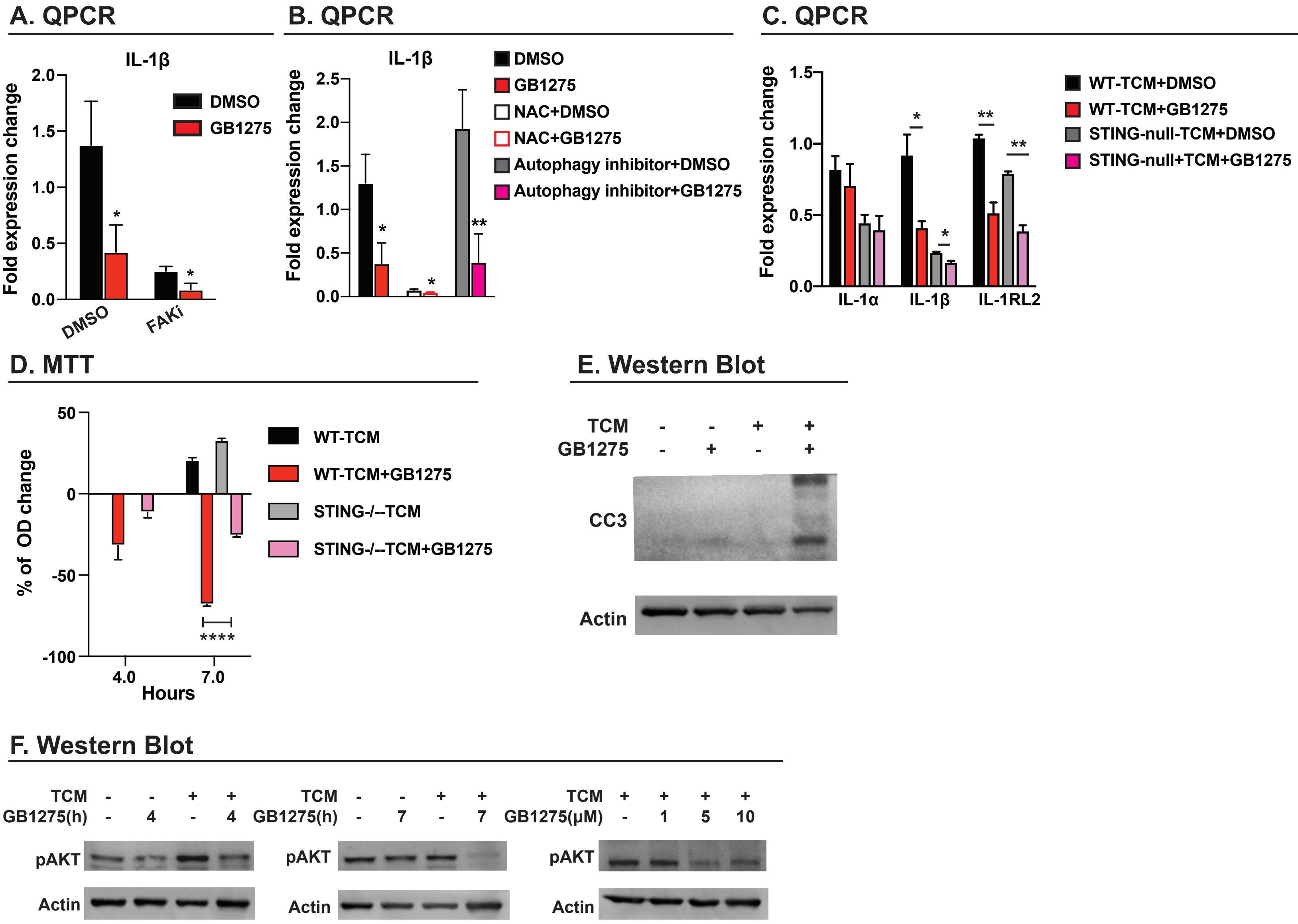

### Supplementary Figure 9

#### A. mpIHC

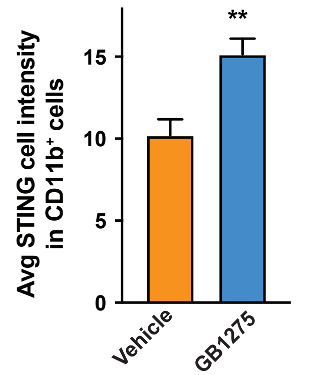


#### B. Human TMA

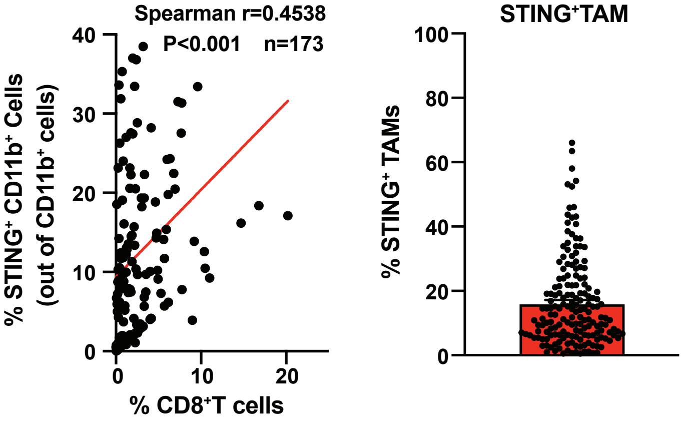


#### C. mpIHC in TMA

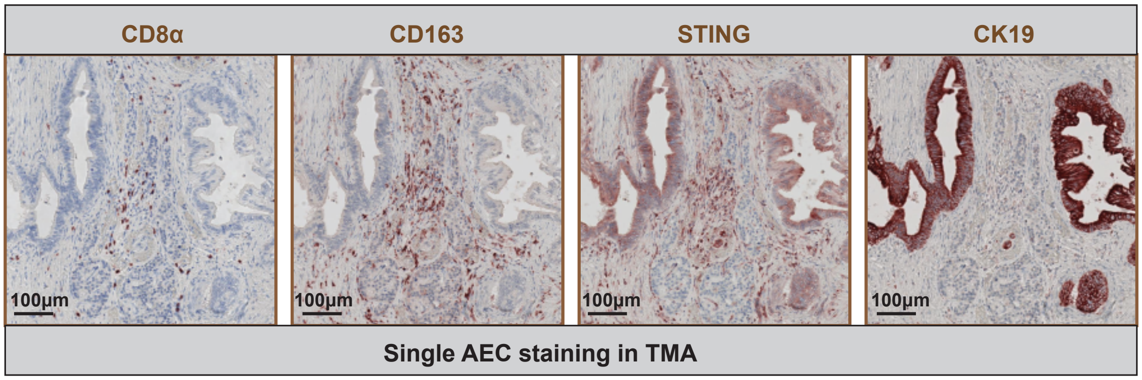


#### D. mpIHC in tumor biopsies

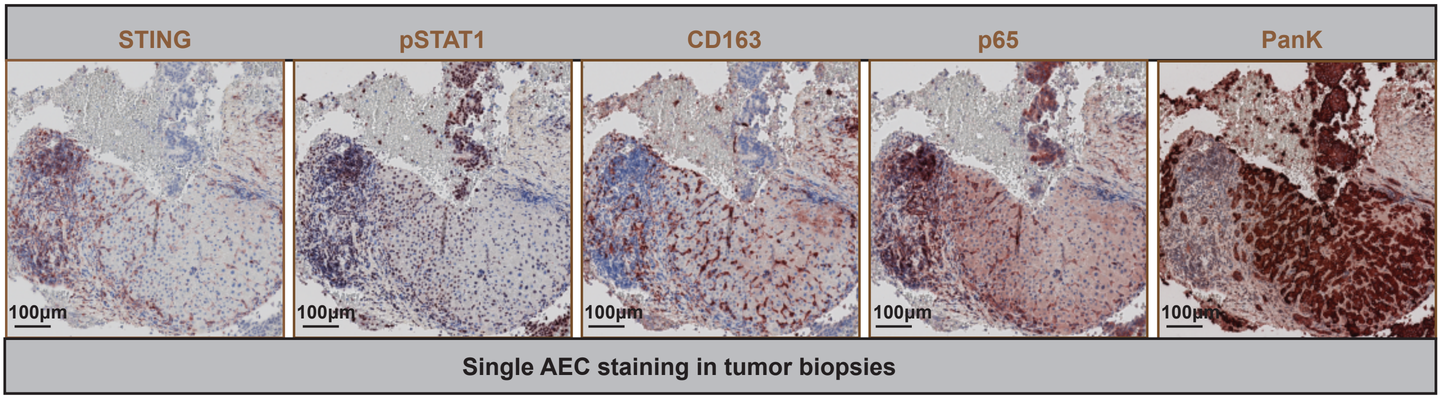


#### E. Tumor biopsies

| Regimen | Tumor Orgin | Number of patient |
|---|---|---|
| A | Colon/Rectum | 4 |
| A | Esophagus | 1 |
| A | Pancreas | 1 |
| B | Colon/Rectum | 2 |
| B | Esophagus | 1 |
| B | Gastric | 1 |
| B | Pancreas | 1 |

### Supplementary Figure 10

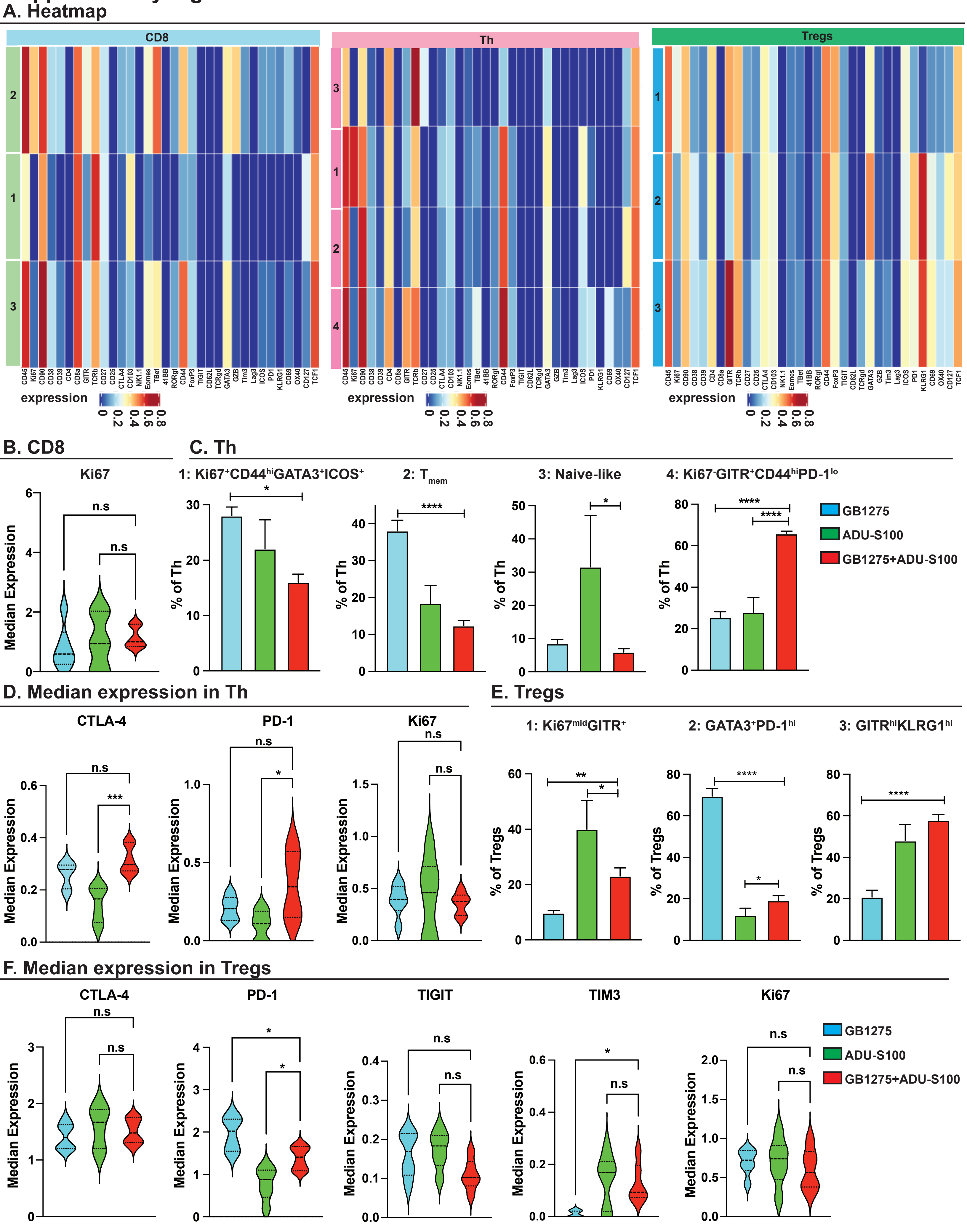

#### Supplementary Figure 11

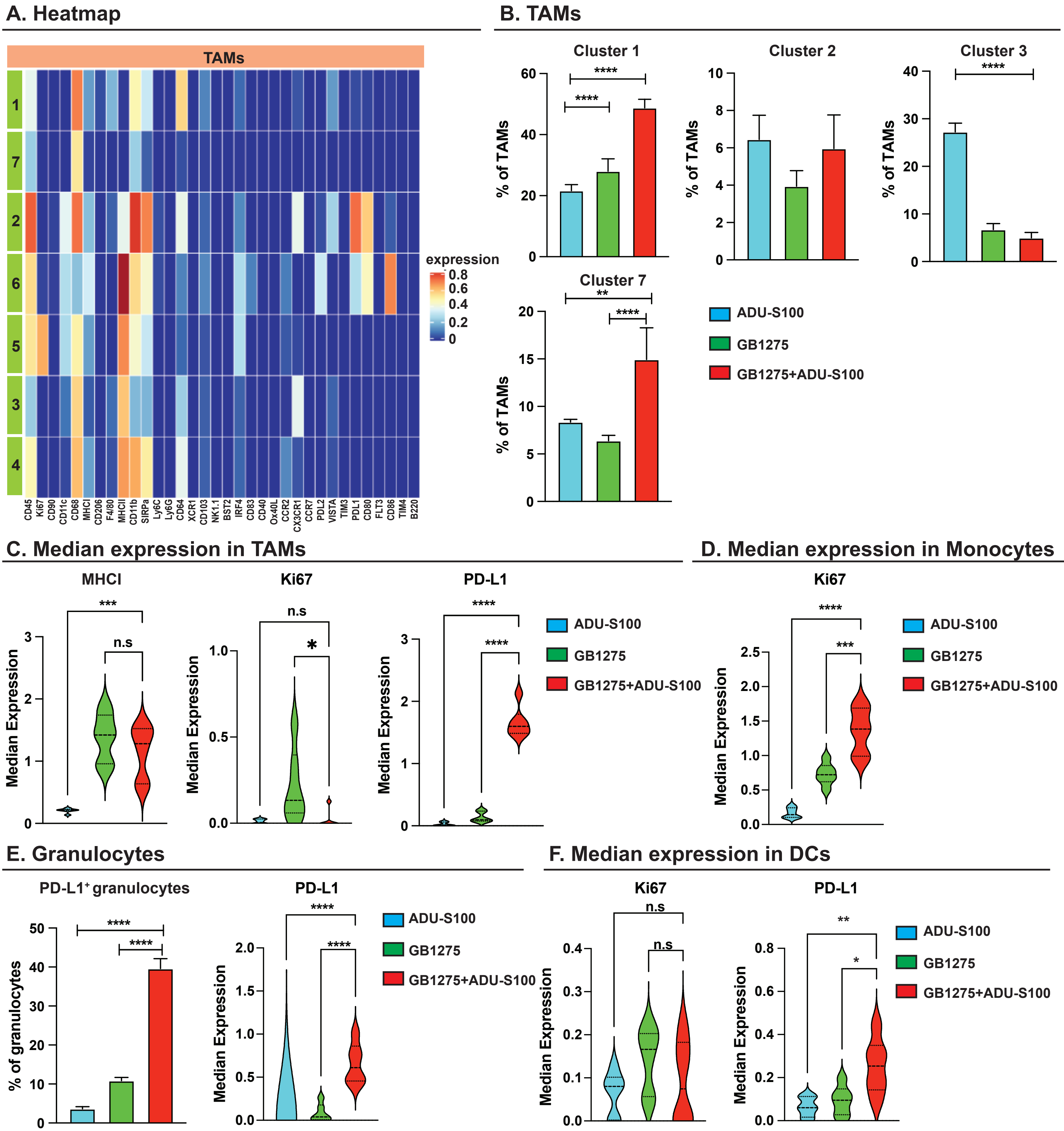

#### Supplementary Figure Legends

##### Supplementary Figure 1 GB1275 exhibits anti-tumor function via CD11b

A. Single UMAP visualization of TAMs (Fig. 1A) shown in each cancer type. Gene set enrichment analysis (GSEA) identified pathway enrichment in TAMs in above cancer types analyzed from curated data sources, including PID **(B)** and BIOCARTA (**C**).

##### Supplementary Figure 2 GB1275 exhibits anti-tumor function via CD11b

**A**. Chemical structures of LA1 and ADH-503 (GB1275). **B**. Syngeneic orthotopic model of KP was built in WT and CD11b- null mice and then treated with Vehicle or GB1275 (120mg/kg) for 14 days (left). Tumor burden from above groups at 14-day treatment shown by tumor volume (n = 6-8/group) (right). **C**. Relative frequencies of tumor infiltrating CD11b$^+$ cells, TAMs, granulocytes and monocytes (n=6/group). **D**. Relative frequencies of tumor infiltrating CD8$^+$ T cells, Ki67$^+$ CD8$^+$T cells, Th cells and the ratio of Th to Tregs (n=6/group). Graphs show the mean ± SEM; * denotes P < 0.05 by two-sided t test or ANOVA test.

##### Supplementary Figure 3 Gate strategies for flow cytometry

**A**. Representative flow cytometry gating strategies for Myeloid cells. **B**. Representative flow cytometry gating strategies for T cells.

##### Supplementary Figure 4 GB1275 affects T cells phenotype

**A**. Heat map of markers stained in CD8$^+$ T cells and CD4$^+$ Th cells and Tregs. **B**. Median Expressions of PD-1, CTLA-4 and TIM3 in CD8$^+$ T cells. **C**. Percentage of individual subclusters in Tregs. **D**. Median expressions of CTLA-4, PD-1 and Ki67 in CD4$^+$ Th cells. **E**. Median expressions of PD-1 and TIGIT and TIM3 in Tregs. Graphs show the mean ± SEM; * denotes P < 0.05 by two-sided t test.

**Supplementary Figure 5 GB1275 inhibits NFκb**

**A**. UMAP scRNAseq plots show $CD45^+$ cells in vehicle and GB1275- treated groups. **B**. Heat map showing differentially expressed genes in subclusters from TAM/monocytes population. **C**. Quantitative PCR mRNA expression analysis of BMDMs treated with Vehicle or GB1275 for 7 hours in the presence or absence of different PDAC conditioned media. Changes in gene expression are depicted as the fold change from the vehicle baseline. **D**. Quantitative PCR mRNA expression analysis of BMDMs co-cultured with KP2 cells ± Vehicle or GB1275 for 48 hours. Changes in gene expression are depicted as the fold change from the vehicle baseline. **E**. Quantitative PCR mRNA expression analysis of human monocytes-derived macrophages treated with different PDAC conditioned media ± Vehicle or GB1275 for 7 hours. Changes in gene expression are depicted as the fold change from the vehicle baseline. **F**. Quantitative PCR mRNA expression analysis of BMDMs isolated from either WT or CD11b- null mice treated with PDAC conditioned media ± Vehicle or GB1275 for 7 hours. Changes in gene expression are depicted as the fold change from the vehicle baseline. Graphs show the mean ± SEM; * denotes $P < 0.05$ by two-sided t test. In vitro data are representative of 3 independent experiments.

**Supplementary Figure 6 GB1275 inhibits inflammatory signaling in macrophages**

**A**. Representative immunoblot for total p65, pP38, pMK2 and β-actin (loading control) in BMDMs treated with GB1275+/- TCM at different doses for indicated time points. **B**. Representative immunoblot for total p65 and β-actin (loading control) in BMDMs transfected with siRNA targeting RelA and ctrl siRNA for 24 h. **C**. Quantitative-PCR mRNA expression analysis of BMDMs transfected with RelA siRNA or ctrl siRNA for 24 hours in prior to 7h- Vehicle or GB1275 treatment. Changes in gene expression are depicted as the fold change from the vehicle baseline. **D**. Quantitative PCR mRNA expression analysis of BMDMs treated with Vehicle or GB1275 for 7 hours. Changes in gene expression are depicted as the fold change from the vehicle baseline. **E**. Percentage of $p65^+$ $CD11b^+$ cells in KPC tumor tissue treated with Vehicle or GB1275 for 14 days (n=6-7 mice per group). **F**. Quantitative PCR mRNA expression analysis of BMDMs transfected with RelA siRNA or ctrl siRNA for 24 hours in prior to 7h- Vehicle or GB1275 treatment with TCM. Changes in gene expression are depicted as the fold change from the vehicle baseline. Graphs show the mean ± SEM; * denotes $P < 0.05$ by two-sided t test or analysis of variance. In vitro data are representative of 3 independent experiments.

**Supplementary Figure 7 GB1275 triggers STING signaling**

**A**. Quantitative PCR mRNA expression analysis of BMDMs isolated from WT and CD11b- null mice treated with Vehicle or GB1275 ± TCM for 7 hours. Changes in gene expression are depicted as the fold change from the vehicle baseline. **B**. Quantitative PCR mRNA expression analysis of BMDMs co-cultured with tumor cells ± Vehicle or GB1275 for 48 hours. Changes in gene expression are depicted as the fold change from the vehicle baseline. **C**. The number of changed proteins following GB1275± TCM at 4h and 7h by RPPA shown in table. **D**. Representative immunoblot for total FAK, pFAK and β-actin (loading control) in BMDMs from WT or CD11b-null mice treated with Vehicle or GB1275 ± TCM for 7 hours. **E**. FAKi (0.5μM)- pretreated BMDMs were stimulated by TCM+GB1275 for 7 hours, and the intracellular level of total ROS was measured by flow cytometry. **F**. Quantitative PCR mRNA expression analysis of NAC (1mM)-pretreated BMDMs treated with TCM ± Vehicle or GB1275 for 7 hours. Changes in gene expression are depicted as the fold change from the vehicle baseline. **G**. Representative immunoblot for STING and β-actin (loading control) in BMDMs transfected with siRNA targeting STING and ctrl siRNA. **H**. Quantitative PCR mRNA expression analysis of BMDMs transfected with STING siRNA or ctrl siRNA for 24 hours in prior to 7h- Vehicle or GB1275 +TCM. Changes in gene expression are depicted as the fold change from the vehicle baseline. **I**. Relative frequencies of tumor infiltrating $Ki67^{+}CD8^{+}$T cells, TAMs, $CD11b^{+}$ cells, granulocytes and eosinophils (n=6/group), and ratio of Th to Tregs in Bone marrow transplant model (BMT) (4K) treated with Vehicle or GB1275 for 14 days. Graphs show the mean ± SEM; * denotes $P < 0.05$

by two-sided t test or analysis of variance. In vitro data are representative of 3 independent experiments.

**Supplementary Figure 8 GB1275 affects macrophages fate**

**A**. Quantitative PCR mRNA expression analysis of FAKi (0.5μM)- pretreated BMDMs treated with Vehicle or GB1275 for 7 hours. Changes in gene expression are depicted as the fold change from the vehicle baseline. **B**. Quantitative PCR mRNA expression analysis of NAC (1mM) or Autophagy inhibitor (10μM)- pretreated BMDMs treated with Vehicle or GB1275 for 7 hours. Changes in gene expression are depicted as the fold change from the vehicle baseline. **C**. Quantitative PCR mRNA expression analysis of BMDMs isolated from WT and STING- null mice treated with TCM ± GB1275 for 7 hours. Changes in gene expression are depicted as the fold change from the vehicle baseline. **D**. MTT proliferation assay using BMDMs treated with TCM ± GB1275 for 4 or 7 hours. Average percentage of OD change after treatment. **E**. Representative immunoblot for cleaved caspase-3 (CC3) and β-actin (loading control) in BMDMs treated with TCM ± GB1275 for 7 hours. **F**. Representative immunoblot for pAKT and β-actin (loading control) in BMDMs treated with different doses of GB1275 for indicated time points in the presence or absence of TCM (same protein and experiment in Sup Fig. 6A and share the same actin). Graphs show the mean ± SEM; * denotes $P < 0.05$ by two-sided t test or analysis of variance. In vitro data are representative of 3 independent experiments.

**Supplementary Figure 9 STING activation in human**

**A**. Average STING intensity in $CD11b^+$ cells from KPC tumor tissue treated with Vehicle or GB1275 for 14 days (n=7-9 mice per group). **B**. Scatter plot showing Spearman's correlation between the percentage of $STING^+$ $CD11b^+$ cells and percentage of $CD8^+$ T cells in human TMA (left). (right) Average percentage of $STING^+$ TAMs from TMA. **C**. Single staining of CD8α, CD163, STING, CK19 in human TMA by mpIHC. Scale bar, 100 μm. **D**. Single staining of STING, pSTAT1,

CD163, p65 and PanK in tumor biopsies by mpIHC. Scale bar, 100 μm. **E**. Information of tumor biopsies, including treatment regimen, cancer type and number of patients. Regimen A (GB1275 monotherapy), Regimen B (GB1275+ pembrolizumab). Graphs show the mean ± SEM; * denotes P < 0.05 by two-sided t test, or log-rank test.

**Supplementary Figure 10 GB1275 combined with ADU-S100 regulates T cells**

**A**. Heat map of markers stained in $CD8^{+}T$ cells, Th and $T^{Regs}$. **B**. Median expression of Ki67 in $CD8^{+}T$ cells. **C**. Percentage of individual subclusters in Th from PDAC tissue treated with GB1275, ADU-S100 or combo group. **D**. Median expressions of CTLA-4, PD-1, Ki67 in Th. **E**. Percentage of individual subclusters in $T^{Regs}$. **F**. Median expressions of CTLA-4, PD-1, TIGIT, TIM3 and Ki67 in $T^{Regs}$. Graphs show the mean ± SEM; * denotes P < 0.05 by analysis of variance.

**Supplementary Figure 11 GB1275 combined with ADU-S100 regulates myeloid cells**

**A**. Heat map of markers stained in TAMs. **B**. Percentage of individual subclusters in TAMs from PDAC tissue treated with GB1275, ADU-S100 or combo group. **C**. Median expressions of MHC-I, Ki67 and PD-L1 in TAMs. **D**. Median expression of Ki67 in Monocytes. **E**. Percentage of $PD\text{-}L1^{+}$ granulocytes and median expression of PD-L1 in granulocytes. **F**. Median expressions of Ki67 and PD-L1 in DCs. Graphs show the mean ± SEM; * denotes P < 0.05 by analysis of variance.

**Supplementary Table 1: Antibody list of T cell panel for mouse mass cytometry time of flight (CyTOF)**

| T cell panel | | |
|---|---|---|
| REAGENT or RESOURCE | SOURCE | Catalog |
| anti-mouse CD44(IM7) | Leinco | C382 |
| anti-mouse GITR(DTA-1) | BioXcell | BE0063 |
| anti-mouse CD25(PC61) | Leinco | C1194 |
| anti-mouse CD38(90) | eBioscience | 14-0381-82 |
| anti-mouse CD90(G7) | Biolegend | 105202 |
| anti-mouse Lag-3(C9B7W) | Leinco | L306 |
| anti-mouse CD27(LG.7F9) | eBioscience | 50-124-94 |
| anti-mouse KLRG1(2F1/KLRG1) | BioXCell | BE0201 |
| anti-mouse CD103(2E7) | Biolegend | 121402 |
| Anti-mouse CD4(GK1.5) | BioXcell | BE0003-1 |
| anti-mouse CD45(30-F11) | Fluidigm | 3089005B |
| anti-mouse CD62L(MEL-14) | Leinco | C2118 |
| anti-mouse ICOS (C398.4A) | eBioscience | 14-9949-82 |
| anti-mouse OX-40(OX-86) | BioXcell | BE0031 |
| anti-mouse PD-1(RMP1-30) | eBioscience | 14-9981-82 |
| anti-mouse TIGIT(1G9) | BioXcell | BE0274 |
| anti-mouse CD69(H1.2F3) | eBioscience | 14-0691-82 |
| anti-mouse TCRb(H57-597) | BioXcell | BE0102 |
| anti-mouse CD127(A7R34) | BioXcell | BE0065 |
| anti-mouse CD39(Duha59) | Biolegend | 143802 |
| anti-mouse NK1.1(PK136) | BioXcell | BE0036 |
| anti-mouse CD8a (53-6.7) | Leinco | C375 |
| anti-mouse TCRgd (GL3) | eBioscience | 14-5711-82 |
| anti-mouse Tim3(RMT3-23) | BioXcell | BE0115 |
| anti-mouse 4-1BB (17B5) | BioLegend | 106107 |
| anti-mouse FoxP3(FJK-16s) | eBioscience | 14-5773-82 |
| anti-mouse GATA3(TWAJ) | eBioscience | 14-9966-82 |
| anti-mouse GranzymeB (GB11) | eBioscience | MA1-80734 |
| anti-mouse CTLA-4(UC10-4B9) | eBioscience | 50-129-16 |
| anti-mouse Ki67(8D5) | Novus | NBP2-22112 |
| anti-mouse TCF1(812145) | R&D | MAB8224 |
| anti-mouse ROR-γт(AFKJS-9) | eBioscience | 14-6988-82 |

| | | |
|---|---|---|
| anti-mouse Eomes (Dan11mag) | eBioscience | 50-245-556 |
| Anti-mouse T-bet(4B10) | Biolegend | 644802 |
| Myeloid panel | | |
| anti-mouse CD45(30-F11) | Fluidigm | 3089005B |
| anti-mouse Ki67(8D5) | Novus | NBP2-22112 |
| anti-mouse CD11C (N418) | Fluidigm | 3142003B |
| anti-mouse CD68(FA-11) | Biolegend | 137001 |
| anti-mouse MHC-I (28-14-8) | Fluidigm | 3144003B |
| anti-mouse CD206 (C068C2) | Fluidigm | 141702 |
| anti-mouse F4/80(BM8) | Fluidigm | 3146008B |
| anti-mouse MHC-II (M5/114.15.2) | Biolegend | 107602 |
| anti-mouse CD11b(M1/70) | Fluidigm | 3148003B |
| Anti-mouse CD172a/SIRPa (P84) | Biolegend | 144002 |
| Anti-mouse Ly6C (HK1.4) | Fluidigm | 3150010B |
| anti-mouse Ly6G(1A8) | Fluidigm | 3151010B |
| anti-mouse CD64(X54-5/7.1) | Fluidigm | 139301 |
| anti-mouse OX-40(OX-86) | BioXcell | BE0031 |
| anti-mouse XCR1(Zet) | Biolegend | 148202 |
| anti-mouse CD103(2E7) | Biolegend | 121402 |
| anti-mouse NK1.1(PK136) | BioXcell | BE0036 |
| anti-mouse Bst2(120G8) | Novous/imgenx | DDX0390P-100 |
| anti-mouse IRF4(3E4) | Fluidigm | 646402 |
| anti-mouse CD39(Duha59) | Biolegend | 143802 |
| anti-mouse NK1.1(PK136) | BioXcell | BE0036 |
| anti-mouse CD83(Michel-17) | thermofisher scientific | 14-0831-82 |
| anti-mouse CD40(HM40-3) | Fluidigm | 124601 |
| anti-mouse Ox40L (RM134L) | Biolegend | 108802 |
| anti-mouse CCR2 (475301) | RnD systems | MAB55381-100 |
| anti-mouse Cx3CR1(SA011F11) | Fluidigm | 3164023B |
| anti-mouse CCR7(4B12) | Fluidigm | 120101 |
| anti-mouse PDL2(TY25) | BioXCell | BE0112 |
| anti-mouse VISTA(MIH63) | Biolegend | 150202 |
| anti-mouse Tim3(RMT3-23) | BioXcell | BE0115 |
| anti-mouse PDL1(10F.9G2) | BioXCell | BE0101 |
| anti-mouse CD80(16-10A1) | Fluidigm | 104702 |

| | | |
|---|---|---|
| anti-mouse CD135/FLT3 (A2F10) | thermofisher scientific | 14-1351-82 |
| Anti-mouse CD86(GL1) | Fluidigm | 3172016B |
| Anti-mouse B220(RA3-682) | Fluidigm | 3144011B |

**Supplementary Table 2: Antibody list of Flow Cytometry**

| Name | Clone# | Fluorophore | Source | Dilution |
|---|---|---|---|---|
| CD45 | 30-F11 | PE-Cy7, BV786 | eBioscience | 1:400 |
| CD3e | 145-2C11 | APC, BV711 | eBioscience | 1:200 |
| CD4 | RM4-4 | FITC | eBioscience | 1:200 |
| CD8a | 53-6.7 | BUV-395 | BD Biosciences | 1:200 |
| Foxp3 | FJK-16s | PE-Cy5 | eBioscience | 1:100 |
| CD19 | eBio1D3 | APC | eBioscience | 1:200 |
| CD11b | M1/70 | Alexa700 | eBioscience | 1:400 |
| Ly6C | HK1.4 | PerCP-Cy5.5 | eBioscience | 1:400 |
| Ly6G | 1A8 | PE | BioLegend | 1:400 |
| F4/80 | BM8 | PE-Cy5 | eBioscience | 1:400 |
| MHCII | M5/115.15.2 | eFluor450 | eBioscience | 1:400 |
| Ki67 | SoADH-5035 | APC | eBioscience | 1:100 |

**Supplementary Table 3: Antibody list for IHC and mpIHC**

| Name | Clone# | Source | Dilution |
|---|---|---|---|
| CD11b | EPR1344 | Abcam | 1:3000 |
| STING | D2P2F | Cell Signaling | 1:500 |
| P65 | D14E12 | Cell Signaling | 1:1000 |
| pSTAT1 | D3B7 | Cell Signaling | 1:200 |
| CK17/19 | D4G2 | Cell Signaling | 1:500 |
| CD8 mouse | D4W2Z | Cell Signaling | 1:200 |
| CD8 human | SP16 | Cell Signaling | 1:100 |
| CD163 | 10D6 | Leica | 1:300 |
| PanK | AE1/AE3 | Leica | Ready to use |

**Supplementary Table 4: Antibody list for western blot and IP**

| Name | Clone# | Source | Dilution |
|---|---|---|---|
| STING | D2P2F | Cell Signaling | 1:1000 |
| P65 | D14E12 | Cell Signaling | 1:1000 |

| | | | |
|---|---|---|---|
| pAKT | D9E | Cell Signaling | 1:1000 |
| pP38 | D3F9 | Cell Signaling | 1:1000 |
| pMK2 | | Cell Signaling | 1:1000 |
| pIRF3 | D6O1M | Cell Signaling | 1:1000 |
| IRF3 | D83B9 | Cell Signaling | 1:1000 |
| pFAK | | Cell Signaling | 1:1000 |
| tFAK | | Cell Signaling | 1:1000 |
| CD8 human | SP16 | Cell Signaling | 1:1000 |
| pP65 | 93H1 | Cell Signaling | 1:1000 |
| pIκB | 14D4 | Cell Signaling | 1:1000 |
| IκB | | Cell Signaling | 1:1000 |
| Cleaved Caspase-3 | Asp175 | Cell Signaling | 1:1000 |
| Ubiquitin | E4I2J | Cell Signaling | 1:500 |
| β-Actin | 13ES | Cell Signaling | 1:4000 |

**Supplementary Table 5: List of qPCR primers**

| Gene | Source | Assay ID |
|---|---|---|
| GAPDH | Taqman | Mm99999915_g1 |
| TBP | Taqman | Mm01277042_m1 |
| HPRT | Taqman | Mm03024075_m1 |
| IFNα1 | Taqman | Mm03030145_gH |
| IFNβ1 | Taqman | Mm00439546_s1 |
| IL-1α | Taqman | Mm00439620_m1 |
| IL-1β | Taqman | Mm00434228_m1 |
| CXCL9 | Taqman | Mm00434946_m1 |
| CXCL10 | Taqman | Mm00445235_m1 |
| CXCL11 | Taqman | Mm00444662_m1 |
| CCL2 | Taqman | Mm00441242_m1 |
| IL-10 | Taqman | Mm00439614_m1 |
| IL-18 | Taqman | Mm00434225_m1 |
| IL-1R1 | Taqman | Mm00434237_m1 |
| IL-1RL2 | Taqman | Mm00519245_m1 |
| mtCOX1 | Taqman | Mm04225243_g1 |
| 18S rRNA | Taqman | Mm03928990_g1 |
| GAPDH (human) | Taqman | Hs02758991_g1 |
| TBP (human) | Taqman | Hs00427620_m1 |

| | | |
|---|---|---|
| HPRT (human) | Taqman | Hs02800695_m1 |
| IL-1β (human) | Taqman | Hs01555410_m1 |

**Supplementary Table 6: Sequences of siRNAs targeting STING (Tmem173) and RelA**

| Clone Name | Sequence |
|---|---|
| mm.Ri.Rela1a.13.1 | 5'-rCrCrUrUrUrArCrUrGrArArArArArGrCrUrArUrUrGrGrACT-3' |
| mm.Ri.Rela1a.13.2 | 5'-rCrArGrUrArUrUrCrCrUrGrGrCrGrArGrArGrArArGrCrACA-3' |
| mm.Ri.Rela1a.13.3 | 5'-rUrArUrGrArGrArCrCrUrUrCrArArGrArGrUrArUrCrArUGA-3' |
| mm.Ri.Tmem173.13.1 | 5'-rGrArArUrCrGrGrGrUrUrUrArUrUrCrCrArArCrArGrCrGTC-3' |
| mm.Ri.Tmem173.13.2 | 5'-rCrUrUrCrUrUrArArUrArArArCrArUrArUrCrUrArUrUrCTC-3' |
| mm.Ri.Tmem173.13.3 | 5'-rArArCrUrUrGrGrArCrUrArCrUrGrUrUrGrArArArArArCCT-3' |
| siNC | na |
